# Multi-regional characterisation of renal cell carcinoma and microenvironment at single cell resolution

**DOI:** 10.1101/2021.11.12.468373

**Authors:** Ruoyan Li, John R. Ferdinand, Kevin W. Loudon, Georgina S. Bowyer, Lira Mamanova, Joana B. Neves, Liam Bolt, Eirini S. Fasouli, Andrew R. J. Lawson, Matthew D. Young, Yvette Hooks, Thomas R. W. Oliver, Timothy M. Butler, James N. Armitage, Tev Aho, Antony C. P. Riddick, Vincent Gnanapragasam, Sarah J. Welsh, Kerstin B. Meyer, Anne Y. Warren, Maxine G. B. Tran, Grant D. Stewart, Sam Behjati, Menna R. Clatworthy, Peter J. Campbell, Sarah A. Teichmann, Thomas J. Mitchell

## Abstract

Tumour behaviour is dependent on the oncogenic properties of cancer cells and their multicellular interactions. These dependencies were examined through 270,000 single cell transcriptomes and 100 micro-dissected whole exomes obtained from 12 patients with kidney tumours. Tissue was sampled from multiple regions of tumour core, tumour-normal interface, normal surrounding tissues, and peripheral blood. We found the principal spatial location of CD8^+^ T cell clonotypes largely defined exhaustion state, with clonotypic heterogeneity not explained by somatic intra-tumoural heterogeneity. *De novo* mutation calling from single cell RNA sequencing data allows us to lineage-trace and infer clonality of cells. We discovered six meta-programmes that distinguish tumour cell function. An epithelial-mesenchymal transition meta-programme, enriched at the tumour-normal interface appears modulated through macrophage expressed *IL1B*, potentially forming a therapeutic target.

**Single sentence summary:** Kidney cancer evolution, prognosis and therapy are revealed by a single cell multi-regional study of the microenvironment.

## Main Text

Clear cell renal cell carcinoma (ccRCC) is the most common subtype of renal cell carcinoma (RCC), accounting for approximately 75% of RCC cases and the majority of deaths from kidney cancer (*1*). Many efforts have characterised the genomic landscape of ccRCC, revealing important driver events such as bi-allelic inactivation of *VHL* (most commonly via concomitant loss of chromosome 3p and mutation/epigenetic silencing in *VHL*), followed by mutations in chromatin remodelling and histone modification related genes *PBRM1*, *BAP1* and *SETD2* (*2–6*). The timing of initiating events in the evolution of ccRCC has been systematically studied (*4*). Intra-tumoural heterogeneity (ITH) of subsequent mutational events appears to be a salient feature of ccRCC, as revealed by previous multiregion exome sequencing studies (*5, 7, 8*). In contrast, the ITH of ccRCC at a transcriptional level is less well understood, in part due to the complexity of the multicellular ecosystem comprising the tumour microenvironment (TME). In particular, the phenotypic heterogeneity of malignant and non-malignant cells in the TME of ccRCC and how it associates with geographical localisation remain elusive.

ccRCC is a cancer type with heavy infiltration of immune cells (*9, 10*). Immune checkpoint blockade (ICB) therapy has been shown effective in improving the survival of patients (*11, 12*), highlighting the importance of exploring the immune microenvironment of ccRCC. Previous studies profiled the immune landscape of ccRCC using bulk sequencing (*9, 13*), which limited their power in dissecting the diverse immune cell population. A comprehensive immune atlas of ccRCC using mass cytometry shed light on immune cell diversity in the ccRCC tumour ecosystem (*14*). Recent advances in single cell RNA sequencing (scRNA-seq) and its applications in cancer research have revolutionised our understanding of phenotypic heterogeneity of tumour cells (*15–17*), immune landscape of tumours (*18–20*), complexity and plasticity of the TME (*21, 22*), and intercellular communications in the TME (*23, 24*). Specifically, in ccRCC, a recent scRNA-seq study provided evidence to support its origin from proximal tubular cells (*25*). Other studies utilised scRNA-seq to study the immune landscape of ccRCC mainly focusing on ICB therapy related cohorts (*26, 27*) and different disease stages (*28*), uncovering key features that are related to therapeutic efficacy or disease progression.

In considering heterogeneity in the TME, the geographic regions of interest extend those relevant to mutational ITH. The wider regions of interest include circulating blood, the tumour-normal interface or tumour pseudocapsule (representing the boundary between tumour and adjacent normal kidney), adjacent normal kidney, and perinephric adipose tissue. The fibrous connective tissue of the pseudocapsule appears to spatially constrain growth, and invasion is correlated with tumour stage and grade (*29*). Perinephric adiposity is of interest because of the obesity paradox in RCC, whereby obesity is one of the strongest risk factors for the diagnosis of kidney cancer, yet is also associated with improved oncological outcomes (*30*). Understanding the spatial heterogeneity and evolution of ccRCC with respect to tumour cells, various immune/stromal cells, and interactions between them in the wider TME is still lacking. To address this, we performed multi-region based scRNA-seq from 12 patients, sampling peripheral blood, normal kidney, four different spatial regions of the tumour core, and the tumour-normal interface, alongside focally exhaustive exome sequencing of laser-capture microdissection (LCM) derived tumour samples.

## Results

### Multi-region based genomic and single-cell transcriptomic profiling of RCC

We conducted multi-region genomic and single-cell transcriptomic profiling in 12 patients, who underwent surgical resection of radiologically diagnosed renal tumours. After histopathological examination, tumours from 10 out of the 12 patients were evaluated as ccRCC, one (PD47172) was an oncocytoma, and one (PD44714) was a large benign thick-walled cyst (fig. S1A and table S1). In each patient, we sampled tissues from peripheral blood, normal kidney, four different spatial regions of the tumour core, and the tumour-normal interface. Additionally, we sampled tissues from the perinephric fat, normal adrenal gland, adrenal metastasis, and tumour thrombus, if available (Fig. 1A). Where sufficient numbers of viable single cells could be retrieved from these samples, we performed droplet-based 5’ scRNA-seq with T-cell receptor (TCR) enrichment using the 10x platform (table S2). In parallel, in each patient, we dissected micro-biopsy samples from each region containing tumour tissue using LCM prior to performing whole-exome sequencing (WES) (table S3). The LCM approach allowed us to explore the limit of ITH by interrogating sub-millimeter sized biopsies. Based on WES data, we identified genomic alterations that have been reported as recurrent/driver events in ccRCC (*2, 4*). Seven out of nine ccRCC patients (no data in one ccRCC patient) harboured *VHL* mutations, four had *PBRM1* mutations, and three carried *BAP1* mutations (fig. S1A and table S1). Copy number loss of chromosome 3p was detected in all of the nine patients (fig. S1A). The oncocytoma carried a characteristic copy number loss of the whole of chromosome 1 (fig. S1A).

**Fig. 1.**
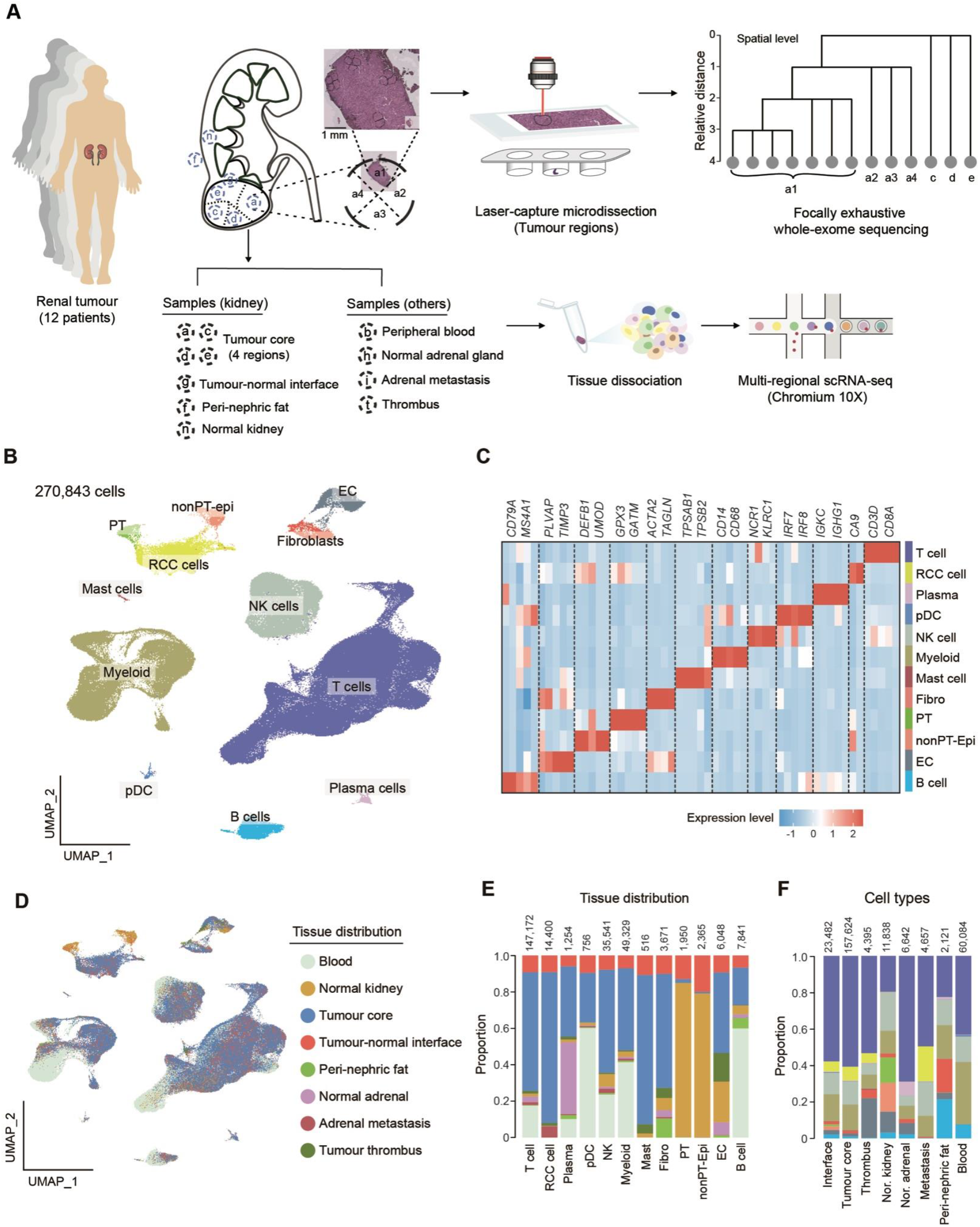
Sampling strategy and overall tissue distribution of the major cell types in RCC. (**A**) Sampling strategy for each of 12 patient donors. a, c, d and e represent four different regions of the tumour core; g, tumour-normal interface; f, perinephric fat; n, normal kidney; b, peripheral blood; h, normal adrenal gland; i, adrenal metastasis; t, thrombus. a1, a2, a3 and a4 represent LCM biopsies in tumour region a. **(B)** Overall uniform manifold approximation and projection (UMAP) of all cells in our study. (**C**) Heatmap showing top differentially expressed genes (DEGs) in each of the major cell types. (**D**) UMAP showing tissue distribution of the major cell types. (**E** and **F**) Barplots showing the tissue distribution of the major cell types. Colours in (E) correspond to those in (D), and colors in (F) correspond to those in (B).

Using scRNA-seq, we captured transcriptomes from approximately 270,000 cells after stringent quality control, which can be broadly categorised into 12 major cell types based on the expression of canonical marker genes (Fig. 1, B and C, fig. S1, B to D). As a result of our single cell isolation protocol, T cells (expressing *CD3D, CD3G,* and *CD3E*) were most abundant in our data, followed by myeloid cells (expressing *CD68* and *CD14*) and natural killer (NK) cells (expressing *NCR1* and *KLRC1*) (Fig. 1C and fig. S1E). We also captured significant numbers of other immune cell populations: B cells (expressing *CD79A* and *MS4A1*), plasma cells (expressing *IGKC* and *IGHG1*), mast cells (expressing *TPSAB1* and *TPSB2*), and plasmacytoid dendritic cells (pDC; expressing *IRF7* and *LILRA4*) (Fig. 1C and fig. S1E). Apart from the immune compartment, four stromal cell types were identified in our data: endothelial cells (EC) (expressing *PLVAP* and *TIMP3*), fibroblasts (expressing *ACTA2* and *TAGLN*), proximal tubular (PT) cells (expressing *SLC22A8* and *GPX3*), and non-PT epithelial cells (expressing *DEFB1* and *UMOD*) (Fig. 1C and fig. S1E). RCC tumour cells were identified within clusters that specifically expressed *CA9* and harboured extensive copy number variations (CNVs) across their genomes, as inferred from scRNA-seq data (Fig. 1C, fig. S1, E and F). Next, we investigated the tissue of origin of the 12 major cell types (Fig. 1D) and observed different tissue distributions of these cell types (four regions were combined in the analysis) (Fig. 1, E and F).

We further conducted sub-clustering analyses for the major cell compartments covered in our study. Sub-clustering of NK cells generated 14 clusters with differentially expressed genes (DEGs) and heterogeneous tissues of origin (fig. S2, A to D and table S4). Among these clusters, the well-known CD56 (*NCAM1*) and CD16 (*FCGR3A*) expressing populations were identified (fig. S2B). An innate lymphoid cell (ILC) cluster was characterised by expression of *IL7R* and *FXYD7* (fig. S2B). Two NK clusters (cluster 2 and 6) showed high expression of interferon gamma (*IFNG*) with cluster 6 also highly expressing cytokine *CCL4L2* and potentially enriching in normal adrenal gland (fig. S2, B and D). Some less characterised NK clusters were identified such as cluster 4, which specifically expressed *KRT81* and *KRT86* (fig. S2B). This NK cell subset was previously reported in hepatocellular carcinomas with its function remaining unclear (*22*). The B/plasma cell compartment was categorised into 13 clusters, among which well-known major B cell populations such as naïve, switched memory, and non-switched memory B cells were identified (fig. S2, E and F, table S4). Activated B cells (both *AREG* and *RHOB* high clusters) were potentially enriched in tumour with *AREG*^*high*^ cluster being more enriched in the perinephric fat. Plasma IgA, IgG and cycling cells (expressing *IGHA1*, *IGHG1* and *MKI67*, respectively) were found to be enriched in tissues compared to peripheral blood (fig. S2, F to H).

### CD8^+^ T cell lineages reveal evolutionary trajectories and the influence of spatial location on exhaustion

T cells comprised the most abundant cell type in our data and were broadly divided into two major compartments: CD4^+^ and CD8^+^ T cells (including gamma delta T cells). Sub-clustering of the CD8^+^ T cell compartment resulted in the identification of 18 clusters with various DEGs and heterogeneous tissue locations (Fig. 2A, fig. S3, A to D). Overall, we identified typical CD8^+^ T cell clusters which represented different T cell functional states including naïve, effector, memory, pre-dysfunction and dysfunction based on the expression of canonical marker genes (Fig. 2A and table S4). Naïve/central memory (CM) CD8^+^ T cells, which highly expressed genes such as *LEF1* and *CCR7*, were found to be mostly enriched in peripheral blood (Fig. 2A and fig. S3D). We identified resident memory (RM) T cells, as they expressed tissue-residency markers (i.e., *ITGAE* and *CD69*) and were mostly enriched in the normal kidney (Fig. 2A, fig. S3, A and D). Particularly, cluster 10 also highly expressed *CXCL13*, which may play potential roles in B cell recruitment and the formation of tertiary lymphoid structures (*31*). We found cluster 6 highly expressed *FGFBP2* and *CX3CR1*, and was substantially enriched in peripheral blood, therefore this cluster may represent recently activated effector memory T cells (CD8^+^ T_EMRA). Two exhausted T cell clusters (cluster 7 and 8) were identified based on elevated expression of genes including *LAG3*, *TIGIT*, *PDCD1*, *HAVCR2*, and *CTLA4* (Fig. 2A and fig. S3A). Interestingly, we found cluster 8 had the highest expression of *LAG3* and specifically expressed the immunosuppressive cytokine *IL10* (Fig. 2A). This cluster may represent CD8^+^ T cells with extremely high effector and dysfunction levels, which exert regulatory functions by producing IL-10. Mucosal-associated invariant T (MAIT) cells were found as highly expressing *TRAV1-2* and *IL7R* (Fig. 2A). Two cycling cell clusters were identified: one (expressing *MCM5* and *PCNA*) represented cells in the G1/S phase of cell cycle and the other (expressing *TOP2A* and *MKI67*) represented those in G2/M phase (Fig. 2A). Besides the conventional CD8^+^ T cell clusters, we also identified two gamma delta T cell clusters: gdT_Vd1 (expressing *TRDV1*) and gdT_Vd2 (expressing *TRDV2*) (Fig. 2A). We also performed sub-clustering analysis of CD4^+^ T cell population, revealing various subtypes such as CD4^+^ naïve/CM and CD4^+^ regulatory T cells (Tregs) and their different tissue distributions (fig. S3, G to I and table S4).

**Fig. 2.**
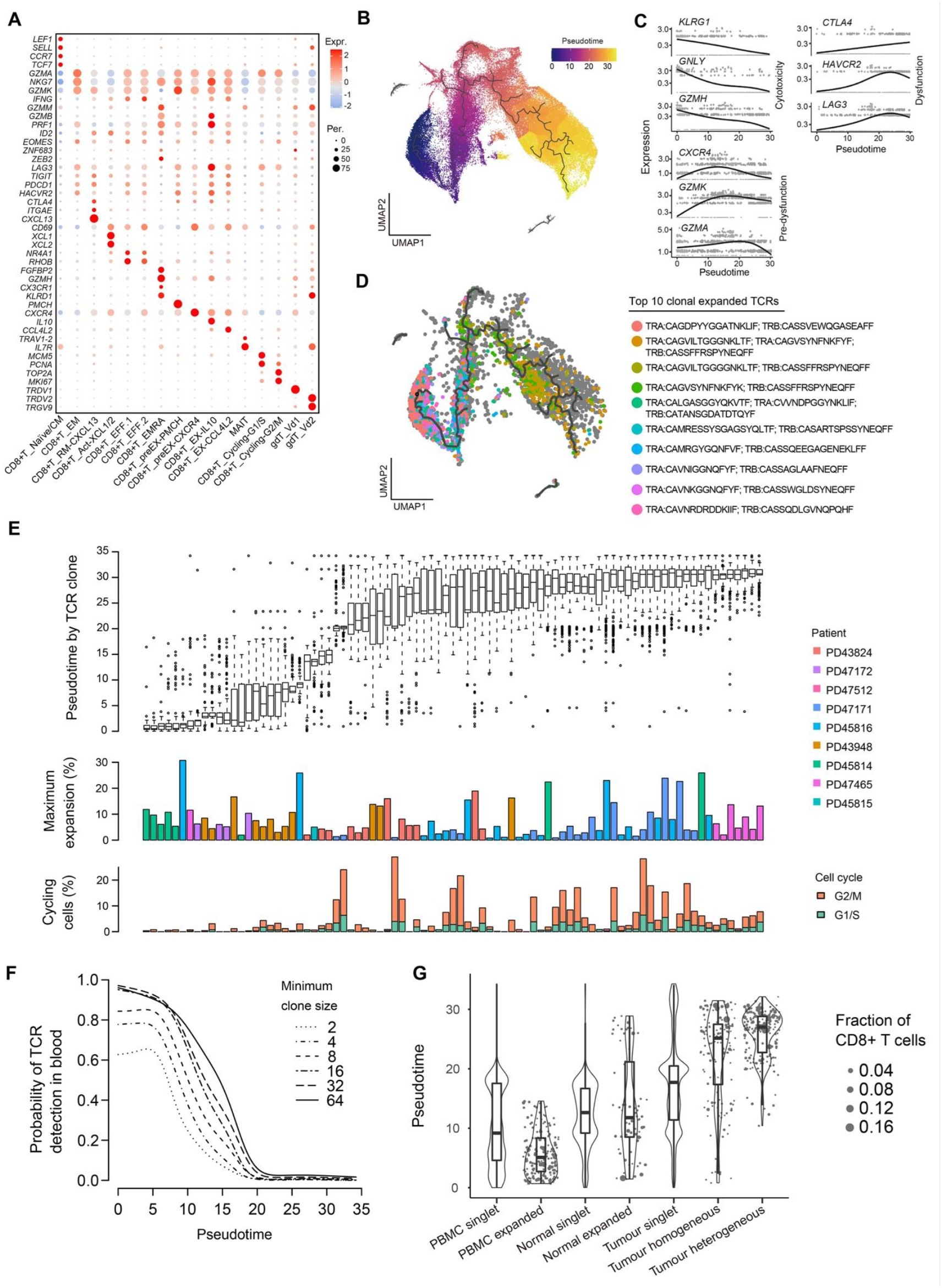
CD8^+^ T cell characterisation, clonality, exhaustion and regional enrichment. (**A**) Dotplot showing marker gene expression defines principal CD8^+^ cell types. EM, effector memory; Act, activated; EFF, effector; EX, exhausted. (**B**) UMAP depicting the pseudotime inference of CD8^+^ cells. **(C)** Expression of canonical exhaustion markers across cells ordered by pseudotime analysis. All marker genes are statistically significant across pseudotime values (*q* value = 0). (**D**) UMAP showing the 10 most expanded clones from patient PD43948. Grey dots represent cells outside the 10 most expanded clones. (**E**) Boxplot depicting the most expanded clonotypes (>100 cells) across all patients, ordered by their mean pseudotime values, showing the median, interquartile range and outlier pseudotime values (top panel); barplots showing the maximum expansion for the most expanded region (middle panel) and percentage of cycling cells (lower panel). (**F**) The probability of detecting a given TCR clone in peripheral blood as a function of minimal clone size and mean pseudotime value of the clone. (**G**) Mean pseudotime values based on the categorisation of clonotypes according to their principal region of enrichment.

Next, we conducted a pseudotime trajectory analysis on CD8^+^ T cells excluding gamma delta T and cycling clusters using the Monocle 3 and RNA velocity analysis (Fig. 2B and fig. S3E). Along the pseudotime trajectory, we found that cytotoxicity related genes (i.e., *KLRG1*, *GNLY* and *GZMH*) were gradually down-regulated while dysfunction related genes (i.e., *CTLA4*, *HAVCR2* and *LAG3*) were gradually up-regulated (Fig. 2C). Typical T cell pre-dysfunction related genes (i.e., *CXCR4*, *GZMK* and *GZMA*) were initially up-regulated and then went down along the pseudotime trajectory (Fig. 2C). Therefore, this pseudotime trajectory recapitulated the progression of CD8^+^ T cells from a cytotoxic state via a pre-dysfunctional state to a dysfunctional state, along with which the degree of exhaustion gradually escalated. This progression was also supported by the positive correlation between the pseudotime and exhaustion score (fig. S3F). Further, projection of the top 10 expanded TCR clonotypes onto the trajectory led to an observation that individual TCR lineages were usually restricted to a similar phenotypic state, rather than distributing across the entire trajectory (Fig. 2D). Across all tumours, we found that 90% of clonotypes with 23 cells or greater were confined within a range of pseudotime values (Wilcoxon test, *p* < 0.05). Highly expanded TCR clones with over 100 cells per clone were observed in multiple patients, where remarkably up to 30% of CD8^+^ T cells can derive from a single clonotype (Fig. 2E). In contrast, TCR clonotypes in CD4^+^ populations were less expanded compared to those in CD8^+^ populations (fig. S3J). Many of the most expanded CD8^+^ TCR clones had considerable proportions of cycling cells, with the exception being observed in the less exhausted clonotypes (Fig. 2E). This finding demonstrates that the proliferation in highly exhausted T cells in RCC has not been completely arrested, similar to previous findings in melanoma (*19*).

We examined whether the TCR clonotypes detected in the blood reflected those detected in other regions. We found the average degree of exhaustion (inferred pseudotime) and the probability of detecting CD8^+^ TCR clones in the peripheral blood were strongly anti-correlated regardless of the clonal size, to the extent that exhausted clonotypes are seldom detected in the blood (Fig. 2F; Method). This finding is unexpected and indicates that tissue-resident exhausted CD8^+^ T cell clones do not appear to recirculate in peripheral blood. To further illustrate the relationship between T cell exhaustion, clonal expansions and their tissue distributions, we categorised CD8^+^ T cells according to whether they were singlets or expanded, and their principal tissue locations (blood, normal tissues or tumour). Expanded T cells in tumour were further subcategorised into those that appeared in all tumour regions and those that do not (tumour homogeneous and heterogeneous). Notably, the phenotypic state of CD8^+^ T cells, in terms of the degree of exhaustion, showed a strong dependence on clonal expansion and tissue location (Fig. 2G; all p < 0.05, Tukey test). Meanwhile, clones that were private to one tumour region were not significantly more exhausted than those shared between different regions (Fig. 2G; p > 0.05, Tukey test).

### Spatial localisation rather than intra-tumoural heterogeneity primarily influences CD8^+^ clonotypic heterogeneity

Using somatic mutations called from WES data, we constructed phylogenetic trees to elucidate the clonal evolution and ITH in tumours in our study. Overall, we found all tumour clones shared a long trunk but had short branches (Fig. 3A). This indicates that most somatic mutations were ubiquitous in individual tumours with only a small number of private mutations being detected. The majority of detected driver mutations and key CNVs (i.e., *VHL* mutations and loss of heterozygosity (LOH) of chromosome 3p) were shared by all tumour clones within individual tumours, thus locating on the trunks of phylogenetic trees (Fig. 3A). Furthermore, the vast majority of LCM samples we sequenced appeared clonal according to the variant allele frequency distributions (fig. S4A). Taken together, the WES revealed that the extent of ITH of tumours in our cohort was limited. Previous studies have extensively investigated intra-tumour genetic heterogeneity in various cancers by comparing somatic mutations detected in samples from different spatial localisations of a tumour (*7, 32*). However, the influence of somatic heterogeneity on the local tumour microenvironment at different spatial localisations, especially the anti-tumour immune response, remains largely uncharacterised. Here we systematically compared the relationship between somatic mutations, spatial localisations and TCR clonotypes of CD8^+^ T cells in individual tumours (Fig. 3B and fig. S4B). Unexpectedly, we found T cell clonotypes were highly heterogeneous and disparate in different spatial localisations, even in tumour regions where only negligible heterogeneity of somatic mutations was observed. Somatic mutations, which generate neoantigens on tumour cells, are considered a driving factor for T cell clonal expansion upon antigen presentation. Our finding suggests the heterogeneity of TCR clonal expansions associate more with the different spatial localisation of T cells in tissues rather than ITH of somatic mutations. To formally examine this, we calculated the correlation between T cell clonotype distance and 1) mutation distance; 2) spatial localisation distance (Fig. 3C). By comparing these two correlations, we found TCR heterogeneity in CD8^+^ T cells was more strongly correlated to spatial localisation rather than somatic heterogeneity (paired Wilcoxon test, *p* < 0.05).

**Fig. 3.**
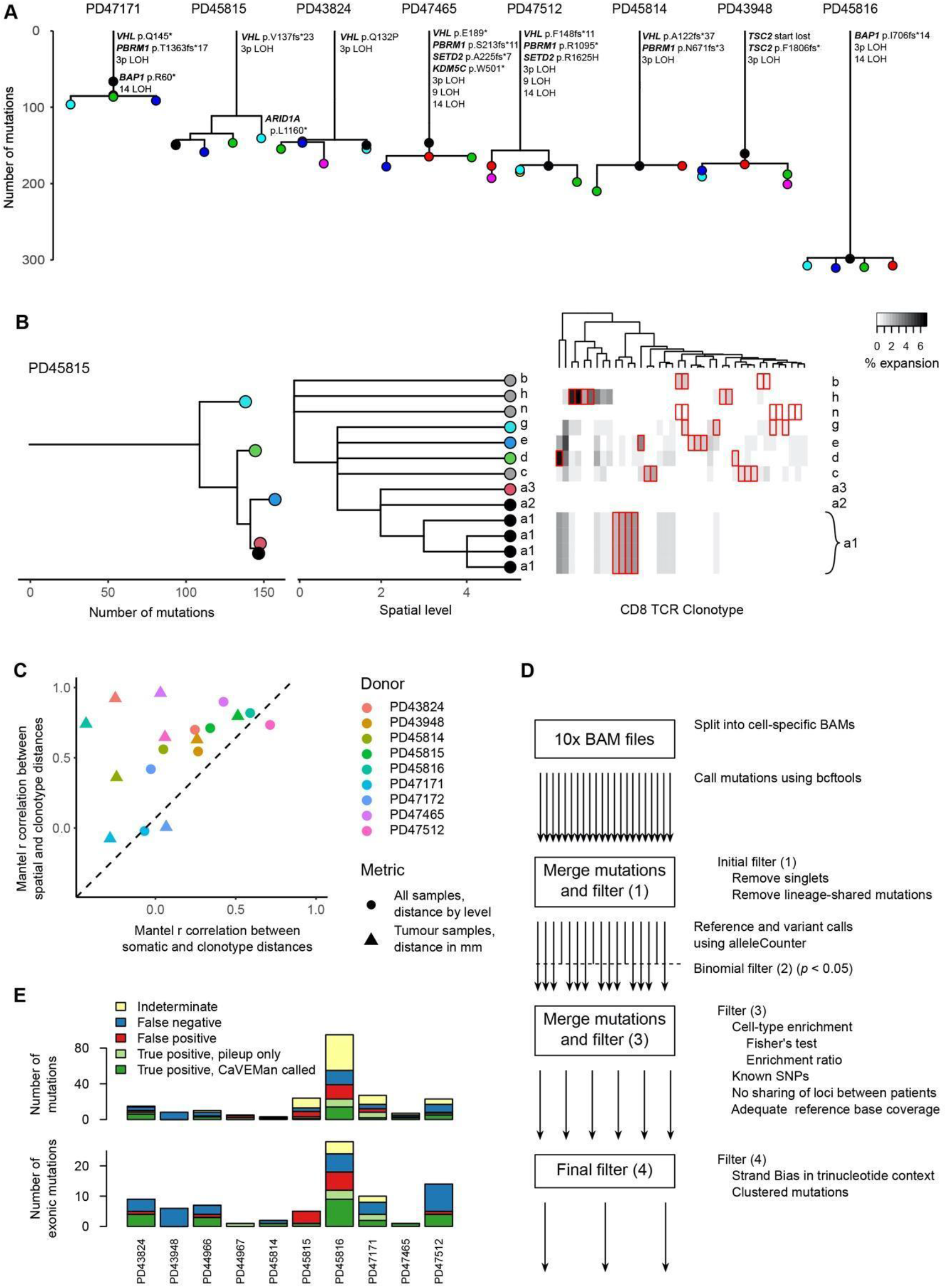
Somatic mutation calling and the relationship with TCR clonotypic heterogeneity. (**A**) Reconstructed phylogenies from WES of multi-regional LCM biopsies. Each node represents a mutant clone present in one or more of the biopsies. (**B**) Comparison of WES derived phylogenies (left) with geographic location (centre) and CD8^+^ TCR clonotype expansion (right). Colours reference somatic clones to spatial localisation. Each column in the right panel represents a TCR clonotype, those with significant regional enrichment are highlighted in red. a, c, d, and e represents four different regions of the tumour core; g, tumour-normal interface; n, normal kidney; b, peripheral blood; h, normal adrenal gland. (**C**) Scatter plot of the Mantel correlation between tree distances. *x-*axis represents the correlation coefficient between WES derived clones and TCR clonotype distances. *y-*axis represents the correlation coefficient between spatial localisation and TCR clonotype distances. (**D**) Schematic of the *de novo* mutation calling algorithm. (**E**) Benchmarking results for scRNA-seq derived calls against WES data for each patient donor.

### Precise *de novo* somatic mutation calling from scRNA-seq data

The detection of somatic mutations within single cells from their transcriptomic sequences may help infer their clonal relationships. Although it is theoretically possible to call mutations from scRNA-seq data, no methods with high accuracy are currently available. Here we developed an algorithm/pipeline to perform *de novo* somatic mutation calling from scRNA-seq data (deSCeRNAMut; Methods). Briefly, we used the bcftools to call mutations from single cell BAM files prior to an initial filter step to remove both mutations present in a single cell and mutations shared between different cell lineages. We checked reference and variant allele counts at all of the loci called in the initial step, prior to applying a binomial filter. We then applied a final set of filtering metrics after annotation of the variants (Fig. 3D). To benchmark our mutation calling method, we first compared somatic mutations called from scRNA-seq data of tumour cells with those called from tumour WES data. All detected mutations were classified as true positive (either detected by the mutation caller, or not called by the mutation caller but with sufficient supporting evidence in the raw data), false positive, false negative, or indeterminate (where mutations were called in regions that did not have sufficient WES coverage to validate the call). Overall, our method achieved a good performance with a precision of 0.64 (or 0.70 when considering exonic mutations only) and a sensitivity of 0.53 (Fig. 3E). We were also able to benchmark the method in CD8^+^ T cells, showing that 84% of called mutations are restricted to a single TCR clone (fig. S4C). This confirms the expected finding that the majority of mutations called in CD8^+^ T cells are restricted to clonotype because of the very limited number of mutations that could be shared between T cell clones pre-thymic maturation.

Using these mutation calls, we investigated the numbers of mutations expressed by different cell types, which can potentially shed light on their degree of clonal expansion. We calculated the proportion of cells with one, two, three, or greater than three mutations. We required at least 100 cells from each cell lineage and patient to account for the lack of discriminatory power in rarer cell populations (fig. S4D). As expected, the lineage with the highest number of cells expressing called mutations were the tumour cells, mainly explained by the known clonal structure of the lineage, but also due to the likelihood of increased mutational burden when compared to the normal cell types. For similar reasons, stromal cells did not typically have discernible numbers of cells with more than one called mutation. However, we observed a surprisingly large number of myeloid cells expressing mutations, indicating that a sizable proportion of these cells are clonally related. These were followed by fibroblasts, and CD8^+^ T cells (which we know are clonally expanded based on the TCR sequencing results). A very small proportion of CD4^+^ T cells expressed mutations, consistent with the low degree of clonality based on TCR analysis (fig. S3J).

### Regional characterisation and evolution of myeloid populations

Overall, we captured transcriptomes from 50,603 myeloid cells in our dataset, which were categorised into 19 clusters in sub-clustering analysis (Fig. 4A, fig. S5A and table S4). We first broadly annotated these clusters based on the expression of canonical markers and the tissue origins of cells (Fig. 4B, fig. S5, B to D). Clusters 1, 2, 3, and 4 were predominantly present in the blood with high expression of *CD14* but lack of *FCGR3A* expression, thus representing circulating classical monocytes. Cluster 5 represented circulating non-classical monocytes with high expression of *FCGR3A* but lack of *CD14* expression (fig. S5D). We identified three dendritic cell (DC) clusters: pDC, type 1 and 2 conventional DC (cDC1 and cDC2), characterised by specific expression of *JCHAIN*, *CLEC9A* and *CD1C*, respectively (fig. S5D). cDC1, among the three DC clusters, showed an enrichment in the tumour core compared with other regions (Fig. 4B). We found mast cells, which were characterised by specific expression of *TPSAB1*, potentially enriched in the tumour core (Fig. 4B and fig. S5D), consistent with previous reports (*33*). Notably, we identified nine macrophage clusters (clusters 6-8, 11-16) based on the high expression of *CD163* and *C1QC* (fig. S5D), reflecting the pronounced heterogeneity of the macrophage population in RCC.

**Fig. 4.**
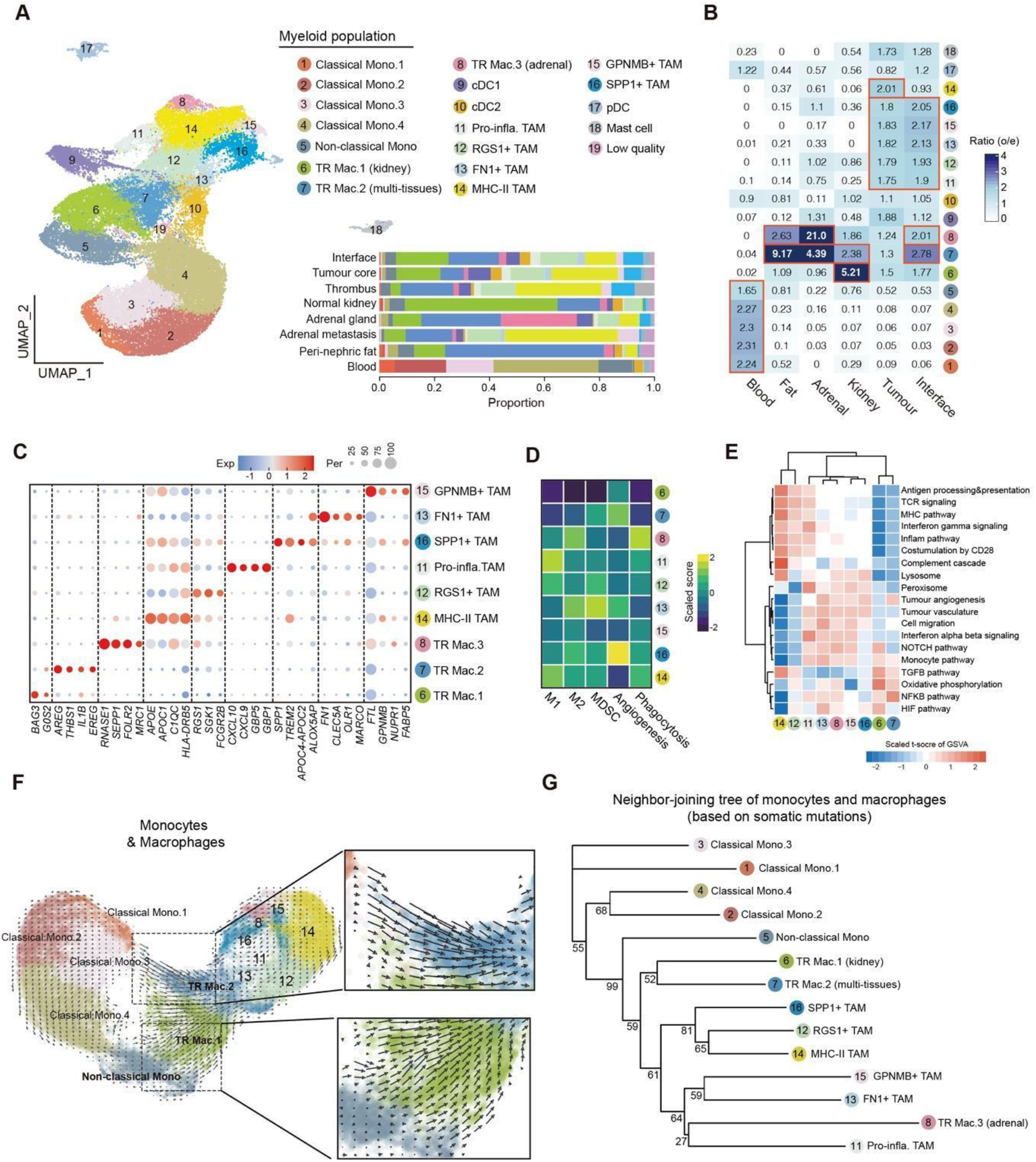
Myeloid cell characterisation, regional enrichment and evolution. (**A**) UMAP representation of all myeloid cells, their annotation and regional contribution. Mono, monocyte; TR Mac, tissue-resident macrophage; TAM, tumour-associated macrophage. (**B**) The relative enrichment of different myeloid cell subsets across different regions sampled. (**C**) Dot plot depicting top DEGs for macrophage clusters. (**D**) Heatmap showing mean scaled scores for macrophage subsets by macrophage function of M1/M2 polarisation, suppressive, angiogenesis and phagocytosis activity. (**E**) Heatmap showing the results of pathway enrichments of macrophage subsets using GSVA analysis. (**F**) UMAP with superimposed RNA velocity analysis of the monocyte and macrophage subsets with zoomed windows highlighting possible directional flows from monocytes to macrophages. (**G**) Neighbour-joining tree depicting the relationship of different monocyte and macrophage clusters, utilising the somatic mutations called from scRNA-seq data. The numbers of supporting votes in bootstrapping (100 times) are labelled.

To further characterise the heterogeneous macrophage population in our dataset, we explored DEGs and the tissue enrichment of the nine macrophage clusters. We found six macrophage clusters (clusters 11-16) preferentially enriched in the tumour core/interface compared to other normal tissues, thus being defined as tumour-associated macrophages (TAMs). The remaining three clusters (clusters 6, 7, 8) showed enrichment in normal tissues and were regarded as tissue-resident macrophages (TR Mac) (Fig. 4B). Among the six TAM clusters, MHC-II TAM (cluster 14) highly expressed *HLA-DRB5*, *APOE* and *APOC1*, and was more enriched in tumour core versus tumour-normal interface. In contrast, the other five TAM clusters showed comparable degrees of enrichments in both tumour core and the interface (Fig. 4B). Pro-inflammatory TAM (cluster 11) highly expressed chemokines *CXCL9/10* and activators of NLRP3 inflammasome assembly *GBP1/5*, exhibiting a predominant feature of M1 polarisation (Fig. 4, C and D, fig. S5E). *FN1^+^* TAM (cluster 15) highly expressed fibronectin 1 (*FN1*) and scavenger receptor *MARCO*, which has been previously reported as a specific macrophage subset in kidney cancer (*33*). We found that *FN1^+^* TAM was likely pro-tumour in ccRCC, as reflected by the high expression of a myeloid-derived suppressor cell (MDSC) signature and of M2 polarization genes (Fig. 4D and fig. S5E). We identified a *SPP1^+^* TAM cluster (cluster 16) in our dataset, which has been reported in various cancer types but as absent in kidney cancer (*33*). Interestingly, we found *SPP1^+^* TAM in ccRCC expressed *GPNMB* and showed a high similarity to the *GPNMB*^+^ TAM identified by the previous study (*33*) (fig. S5F). Considering we also identified a *GPNMB*^+^ TAM cluster (cluster 15) and the expression of *GPNMB* can be detected in multiple TAM clusters (Fig. 4C and fig. S5G), this finding suggests *SPP1^+^* TAM may represent a subset of *GPNMB*^+^ TAM. Besides expressing *SPP1*, we found *SPP1^+^* TAM also expressed *TREM2* and harboured a high angiogenesis score (Fig. 4, C and D). *TREM2^+^* macrophages have been implicated in various biological and pathological processes, such as obesity and cancer (*34, 35*).

Among the three TR Mac clusters, TR Mac.2 highly expressed interleukin *IL1B* and the epidermal growth factor receptor ligand *AREG*, which may reflect its likely role in tissue repair in homeostasis (Fig. 4C). TR Mac.3, which showed high expression of *SEPP1* and *MRC1* and was extremely enriched in normal adrenal gland (Fig. 4, B and C). Interestingly, TR Mac.3 exhibited extremely high expression of M2 and phagocytotic signatures, and showed similar pathway activations to the pro-tumour TAM clusters (i.e., *FN1^+^* TAM) (Fig. 4, D and E, fig. S5E). We were not able to clearly separate embryologically seeded versus monocyte derived tissue macrophages in this dataset (*36*).

Next, we explored the potential origin of different TR Mac and TAM clusters identified in our study. Using RNA velocity analysis, we found two obvious directional flows from circulating monocytes to macrophages in the tissue: (1) classical mono.3 to TR Mac.2 and (2) non-classical monocytes toward TR Mac.1 (Fig. 4F). TR Mac.1 and TR Mac.2 then potentially gave rise to other macrophages in the tissues (Fig. 4F). To determine how macrophage subsets were related to circulating monocytes, we leveraged the somatic mutations for lineage tracing, in a similar way to how the relationship of T cell phenotypic states have been determined from the sharing of TCR clonotypes. Here, we constructed a neighbour-joining tree to depict the relationship of different monocyte and macrophage clusters, utilising the somatic mutations called from scRNA-seq data (Methods; Fig. 4G). We found circulating monocytes were separate from macrophages in tissues and non-classical monocytes (cluster 5) showed a closer relationship with macrophages in tissues compared to other classical monocytes. Our data supports non-classical monocytes representing an intermediary state between circulating monocytes and macrophages, with the majority of macrophages appearing to arise from monocyte progenitor rather than yolk sac origin.

### Regional heterogeneity of endothelial cells, fibroblasts and epithelial cells

We observed heterogeneous stromal cell populations in our dataset. Sub-clustering of the endothelial cell (EC) compartment revealed 11 clusters with different DEGs and preference in tissue locations (Fig. 5, A to C and table S4). Pericytes (cluster 11), as characterised by the expression of *RGS5* and *TAGLN*, preferentially enriched in the tumour core. A small cluster (cluster 9) was found to be extremely enriched in the perinephric fat and highly express *TFF3* and *PDPN*, thus representing lymphatic EC. The remaining nine clusters represented vascular ECs, among which cluster 10 represented cycling EC as highly expressing *TOP2A* and *MKI67*. We identified three potential tumour associated EC clusters: collagen EC, *IGFBP3^+^* EC and *ACKR1*^+^ EC, as they showed considerable enrichments in tumour tissues (Fig. 5, A and C). Among the three clusters, collagen EC (expressing *COL4A1* and *COL15A1*) was found to be more enriched in the interface, which may play roles in interacting with other cells through extracellular matrix (ECM) production. *ACKR1*^+^ EC specifically expressed atypical chemokine receptor ACKR1 which supports adhesion and tissue migration of immune cells (*37*). We found *IGFBP3^+^* EC also expressed a high level of immunosuppressive enzyme *IDO1*, implying its immune regulation roles in the TME. We identified *CRHBP*^+^ EC and *IGF2*^+^ EC preferentially enriching in the normal kidney tissues, while *DNASEL3*^+^ EC showed an enrichment in the normal adrenal gland (Fig. 5C).

**Fig 5.**
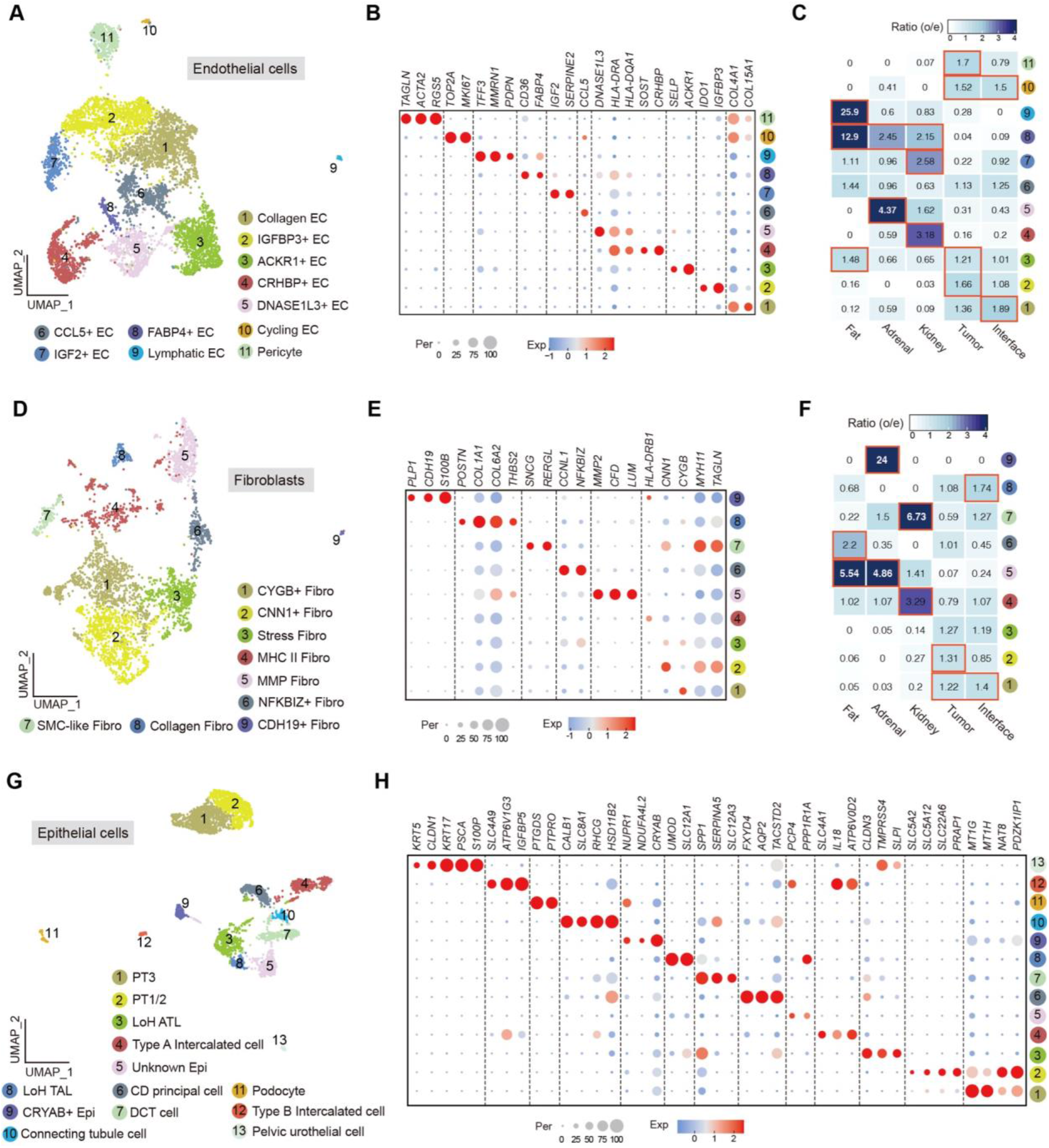
Spatial and transcriptomic heterogeneity of endothelial cells, fibroblasts and epithelial cells. (**A**) UMAP showing the sub-clustering result, (**B**) dot plot depicting top DEGs, and (**C**) regional enrichment of cell clusters of all endothelial cells. (**D**) UMAP showing the sub-clustering result, (**E**) dot plot depicting top DEGs, and (**F**) regional enrichment of all fibroblasts. (**G**) UMAP showing the sub-clustering result and (**H**) dot plot depicting top DEGs of all epithelial cells.

We identified nine clusters of fibroblasts (Fibro) in the sub-clustering analysis (Fig. 5D and table S4). Similar to EC, we found a cluster of fibroblasts (cluster 8) highly expressed collagen-related genes (*COL1A1* and *COL6A2*) and preferentially enriched in the interface (Fig. 5, E and F). This suggests that different ECM-producing stromal cells tend to enrich and co-localise in the interface, exerting diverse functions including extracellular context remodeling and cell-cell interactions. MMP fibro was characterised by high expression of matrix metalloproteinase *MMP2*, complement factor *CFD* and lumican *LUM.* MMP fibro was found to be enriched in perinephric fat and adrenal gland (Fig. 5F). Cluster 7 highly expressed *MYH11*, *SNCG* and *RERGL*, thus being considered as smooth muscle cell (SMC) like fibroblasts. SMC-like fibro was found to be enriched in the normal kidney tissue (Fig. 5F).

The normal epithelial cell population in our dataset exhibited expected diversity (*38*) and was categorized into 13 clusters in the sub-clustering analysis (Fig. 5, G and H, table S4). We identified two proximal tubular (PT) cell clusters (both expressing *NAT8* and *PDZK1IP1*): cluster 1 had higher expression of metallothionein *MT1H* and *MT1G*, thus may representing PT3 cells, while cluster 2 may represent PT1/2 cells as it showed higher expression of PT2 marker *SLC22A6*, and PT1 markers *SLC5A12* and *SLC5A2* (Fig. 5H and table S4). Two Loop of Henle (LoH) clusters: cluster 3 represented ascending thin limb (ATL) cells as expressing *CLDN3* and *TACSTD2*; cluster 8 represented thick ascending limb (TAL) cells, as characterised by expression of *SLC12A1* and *UMOD* (Fig. 5H and table S4). Three collecting duct (CD) epithelial cell clusters were identified: type A intercalated cells (expressing *SLC4A1*), type B intercalated cells (expressing *SLC4A9*) and CD principal cells (expressing *AQP2* and *FXYD4*). We identified distal convoluted tubule (DCT) cells (cluster 7) specifically expressing *SLC12A3*, connecting tubule cells (cluster 10) highly expressing *SLC8A1* and *CALB1*, pelvic urothelial cells (cluster 13) showing specific expression of *PSCA* and *KRT17*, and podocytes (cluster 11) exclusively expressing *PTGDS* and *PTPRO* (Fig. 5H and table S4).

### RCC expression meta-programmes show differential abundance at the tumour-normal interface and affects prognosis

To explore the intra-tumour expression heterogeneity in the tumour cell population, we first defined intra-tumour expression programmes that consist of co-expressed genes in each tumour using non-negative matrix factorization (NMF) (Methods). These expression programmes represented gene modules that were highly expressed by only subsets of tumour cells in each tumour, as exemplified by the NMF result in a representative tumour PD45816 (Fig. 6A). In total, we dissected 45 intra-tumour expression programmes from the ten ccRCC tumours (table S5). Some of these programmes, although subpopulation events in individual tumours, were found to be shared by different tumours, thus being defined as meta-programmes expressed by tumour cells in ccRCC. Six meta-programmes were identified through clustering analysis (Fig. 6B and table S5). The first meta-programme (MP1) was characterised by expression of genes such as *FOS* and *JUN*, thus representing a stress response related signature in tumour cells. MP2 consisted of genes (i.e., *NAT8* and *ACSM2B*) that were specifically expressed by proximal tubular (PT) cells. The presence of PT signature among tumour cells confirmed the previous finding that PT cells are the cell type of origin of ccRCC (*25*). Interestingly, we found a third meta-programme (MP3) was enriched for genes such as *TGFBI* and *MT2A* (Fig. 6B and table S5), which are related to the epithelial-to-mesenchymal transition (EMT). This indicates that MP3 may recapitulate the EMT process in ccRCC, which has not been reported in previous scRNA-seq studies of RCC (*26–28*). MP4 consisted of noncoding RNA genes like *NEAT1* and *HCG18*, probably reflecting some stress or cell death (CD) related cell state. MP5 was characterised by expression of MHC-II related genes such as *CD74* and *HLA-DRA*. Genes such as *TOP2A* and *MKI67* were found in MP6, indicating this meta-programme is related to the proliferation of tumour cells.

**Fig 6.**
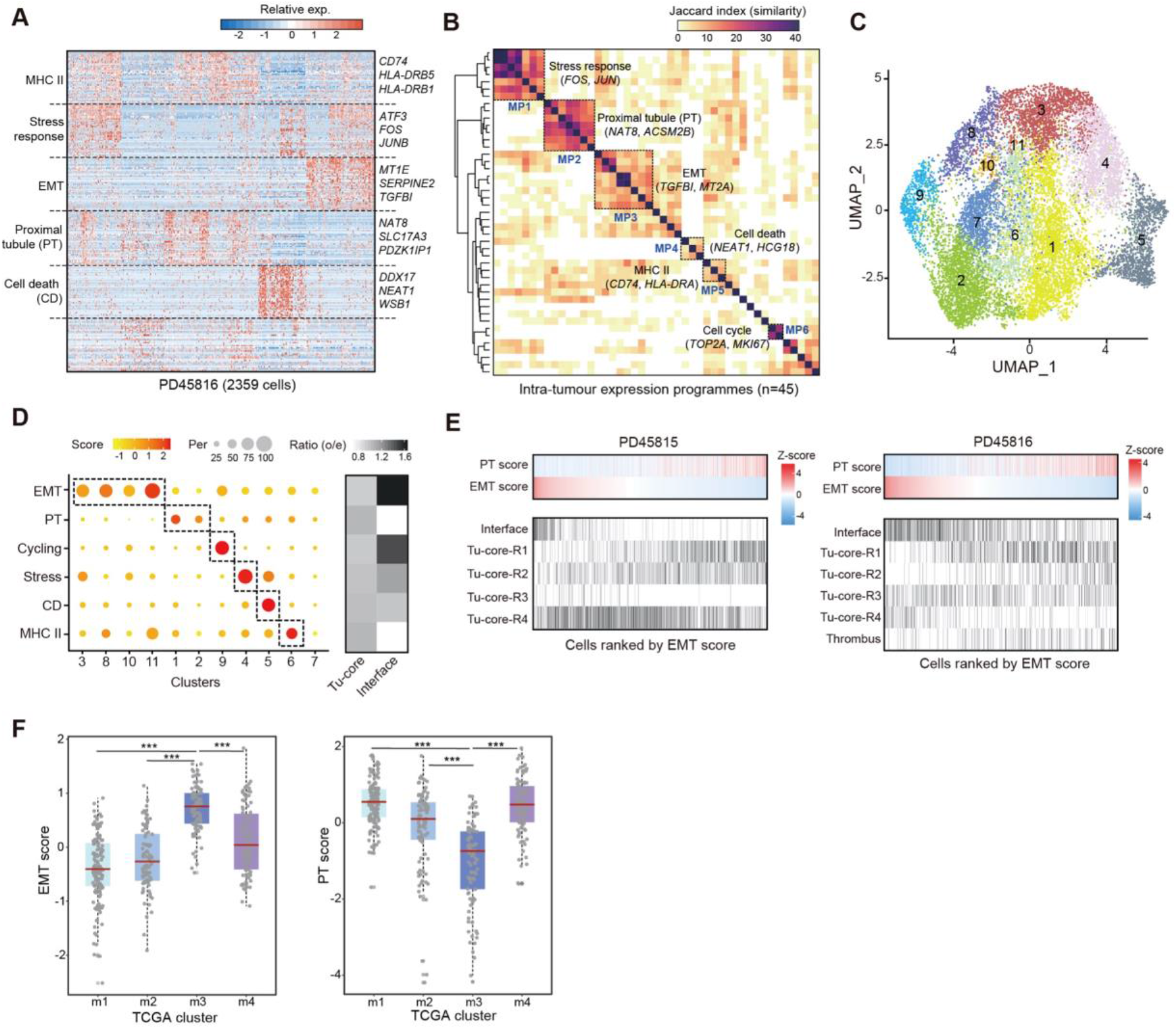
RCC cell expression programmes, regional enrichment and prognosis. (**A**) Heatmap showing expression programmes derived in a representative patient using NMF. (**B**) Heatmap depicting shared expression meta-programmes across all patients. (**C**) UMAP representing clusters of tumour cell population. (**D**) Relative expression scores of meta-programmes in each RCC cell cluster (left) and the distributions of cells with different meta-programmes in tumour core and tumour-normal interface. (**E**) Cells from patient donors PD45815 and PD45816, ranked by decreasing EMT score with corresponding PT score and cell location. (**F**) Box plots showing the EMT and PT scores of TCGA samples in different molecular subtypes. ***, P < 0.001 (Wilcoxon rank sum test).

Next, we integrated tumour cells from the ten tumours, mitigating the inter-patient heterogeneity through batch effect removal (Fig. 6C, fig. S6A, and table S4). Through sub-clustering and DEG analysis, we validated the presence of the six meta-programmes among tumour cells (fig. S6B). We calculated gene scores of the six meta-programmes deciphered using NMF and mapped them onto the UMAP of tumour cells (fig. S6C). This again reflected the expression of the meta-programmes that were sub-populational among tumour cells. Interestingly, we found the expression of PT and EMT programmes showed an inverted pattern (fig. S6C), which was further confirmed by the anti-correlation between PT and EMT scores calculated in bulk RNA-seq data of TCGA samples (fig. S6D). Further, we found that EMT^high^ tumour cells were more abundant at the tumour-normal interface (the leading edge of a tumour) compared to the tumour core (Fig. 6D), which reflects the fact that the EMT state represents a more invasive and migratory state of tumour cells. The heterogeneous expression of PT/EMT programmes coupling the spatial location preference of tumour cells was exemplified by individual tumours (Fig. 6E and fig. S6E). For example, in tumour PD45815, we found PT programme was inversely expressed when ranking tumour cells by EMT scores. Meanwhile, EMT^high^ tumour cells were inclined to locate at the interface while PT^high^ cells were relatively more enriched at region R1 and R2 of the tumour core (Fig. 6E). Finally, by scoring bulk RNA-seq data of TCGA samples, we found that the TCGA molecular subtype m3, which displays the worst prognosis according to TCGA study (*2*), showed significantly higher EMT scores but lower PT scores compared to other subtypes (Fig. 6F). This finding indicates that our meta-programmes can be potential indicators of the survival of patients (fig. S6F).

### Cellular interactions associated with spatial location reveal biological insights and promising therapeutic targets

To characterise intercellular communications in different spatial locations in the RCC microenvironment, we assessed cell–cell interactions between the major cell types in the normal kidney, tumour-normal interface and tumour core using CellPhoneDB (*39, 40*). Interestingly, the numbers of cell–cell interactions occurring among the 12 major cell types in the interface and the tumour core were relatively comparable, which was about two-fold greater than that in the normal kidney tissue even when we excluded those interactions involving tumour cells (Fig. 7A). Next, we compared cell–cell interactions mediated by specific ligand–receptor pairs expressed on different cell types in the normal kidney, tumour-normal interface and tumour core (Fig. 7B). First, this analysis revealed remarkable differences in interactions between the tumour and normal microenvironment. For example, between CD8^+^ T cells and ECs, we found the EC recruitment (*CCL5*–*ACKR1*) and immune inhibitory (*LGALS9*–*HAVCR2*) signals were more active in the interface and tumour core, reflecting escalated levels of angiogenesis and immunosuppression in the tumour compared to the normal tissue (Fig. 7B). Second, we consistently observed differences between the edge and the core of the tumour. For example, a potentially immunosuppressive interaction mediated by *PVR*–*TIGIT* between ECs and CD8^+^ T cells respectively was more active in the tumour core, while a cell growth/migration-related interaction, *IGF*–*IGF2R,* was more active in the interface (Fig. 7B). Third, tumour cells at the interface expressed transcripts predicted to mediate unique intercellular interactions that may potentially promote tumorigenesis. For example, in the interface, myeloid cells express an enhanced cell migration-related signal that can interact with tumour cells (*THBS1*–integrin *α3β1*) compared to that in the tumour core (Fig. 7B).

**Fig 7.**
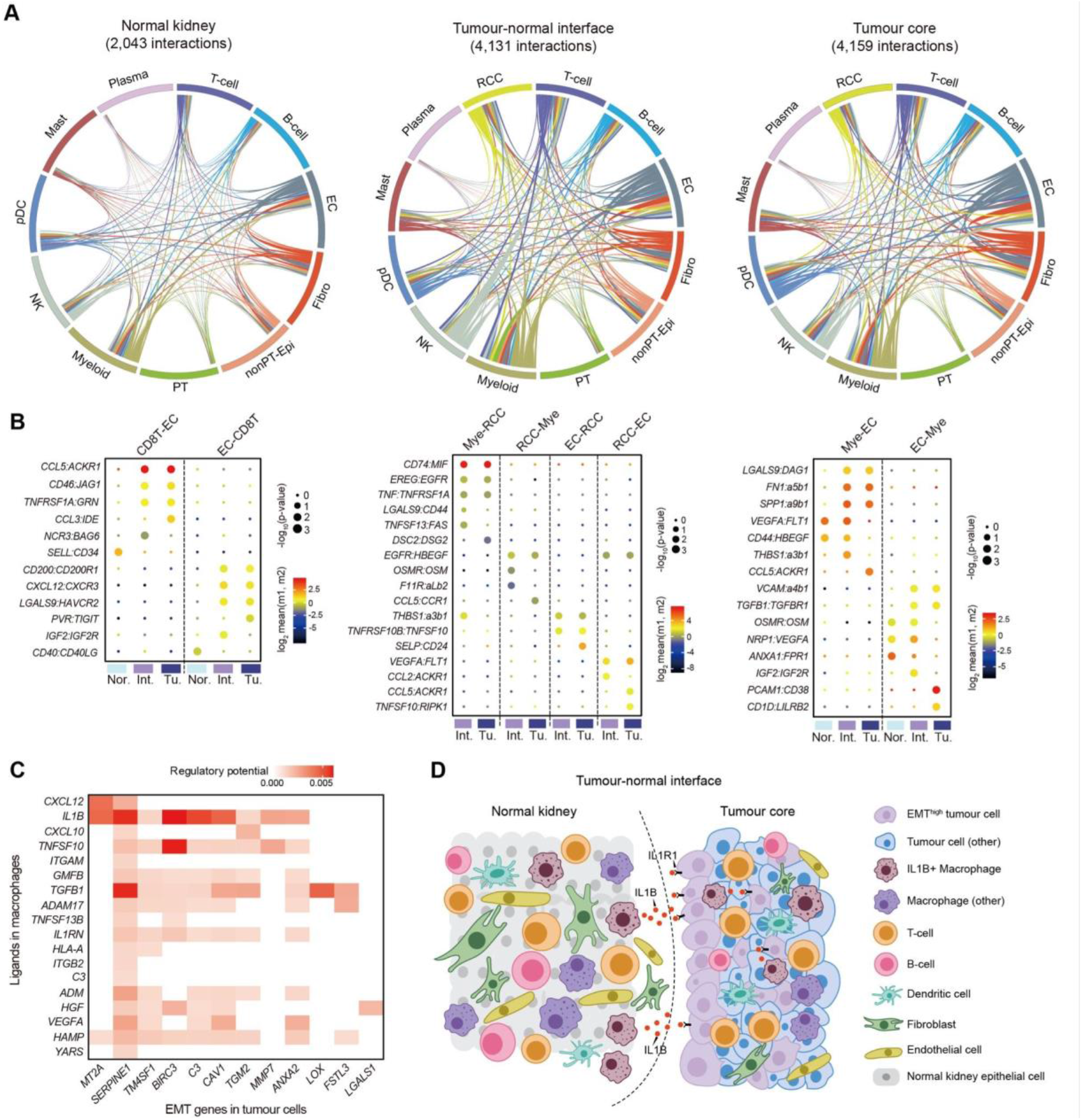
Cellular interactions in the ccRCC micro-environment. (**A**) Comparison of the number of cell– cell interactions between cells present in the adjacent normal kidney, tumour-normal interface, and the tumour core. (**B**) Dot plots representing mean expression levels and significance of important differentially expressed ligand receptor pairs by spatial region. (**C**) Heatmap depicting the potential regulation of genes expressed by the EMT meta-programme and ligands expressed by macrophages. (**D**) Schematic illustrating the importance of IL1B signalling between IL1B^+^ macrophages and RCC cells in promoting EMT.

Our results indicated that tumour cells with high expression of EMT signature (EMT^high^ tumour cells) preferentially localised to the leading edge of tumour (Fig. 6, D and E). This prompted us to explore if there were any active intercellular interactions at the interface that potentially promoted EMT in tumour cells. We used the NicheNet (*41*) analysis to link ligands from cells in the TME and the EMT programme in tumour cells. From this analysis, we found ligands expressed by macrophages potentially regulated a substantial set of EMT genes expressing on tumour cells (Fig. 7C). Particularly, macrophage-derived *IL1B* showed a high and wide regulatory potential to these EMT genes (Fig. 7C), possibly via the receptor *IL1R1* expressed in tumour cells. Interestingly, we found that *IL1B* was specifically expressed by TR Mac.2 (Fig. 4C), which again was preferentially enriched at the tumour-normal interface (Fig. 4B). Taken together, our findings indicate that *IL1B* expressing macrophages (TR Mac.2) preferentially residing at the tumour-normal interface up-regulate the EMT programme in tumour cells at the leading edge of tumours through producing IL1B. Such an oncogenic pathway may eventually promote the migration and invasion of tumour cells (Fig. 7D).

## DISCUSSION

We used multi-region based genomic and single-cell transcriptomic sequencing to characterise the phenotypic heterogeneity and the multicellular ecosystem of ccRCC. Overall, our study depicts a comprehensive atlas of the TME of ccRCC alongside the established ITH in ccRCC, including the phenotypic categorisation of tumour cells and immune/stromal cells, and their intercellular communications in the TME, largely associating with their geographical localisation.

Cells within expanded CD8^+^ TCR clonotypes were largely restricted by exhaustion score. Similar observations were recently reported in melanoma (*42*). The phenotypic restriction of clonotypes may be related primarily to temporal maturation of given clones, rather than environmental factors as individual tumours harboured clonotypes across the full diversity of states. In addition to the phenotypic restriction, we also found that expanded TCR clonotypes were also frequently spatially restricted within one or more of the macroscopic tumour biopsies. This spatial restriction of TCR clonal expansion cannot be entirely accounted for through the exposure to different mutation-associated neoantigens because of the limited ITH of somatic mutations observed in our study. We were unable to define any other factors within the TME that could predict this clonotypic spatial heterogeneity. The perceived stochastic localisation of T cell clonotypes may be a result of physical and environmental factors driving the initial migration of the cells from peripheral circulation to tumour residence. Longitudinal sampling strategies or methods to determine accurate T cell phylogenies could be employed to interrogate the precise timing of T cell expansion and migration. Whether our observation of regional restriction of expanded TCR clones is found more widely in other cancer types may require additional studies deploying similar sampling and sequencing strategies.

The utility of peripheral TCRs for non-invasive cancer detection and surveillance shows promise (*43*), especially in RCC where circulating tumour DNA fragments are scarce (*44*). Although we found many expanded clonotypes were represented both in blood and tumour regions, we observed that the degree of exhaustion and the probability of detecting TCR clones in the peripheral blood were inversely correlated, to the extent that exhausted clonotypes are seldom detected in the blood (Fig. 2F). This finding suggests that once T cell clones infiltrate into tumours and undergo phenotypic transition from activation to dysfunction they seldomly recirculate, possibly due to a tissue residency phenotype as evidenced by CD69 (Fig. 2A). Peripheral sampling of tumour-reactive TCRs is therefore more likely to detect antecedents of exhausted tumour-resident clones, rather than those currently active in the tumour.

We developed a strategy (deSCeRNAMut) to accurately detect somatic mutations in different cell populations based on droplet-based scRNA-seq data. The principal challenges of the lack of consistent coverage, low read depth, and error-prone sequencing reads, were abrogated using a number of filtering metrics including the implausibility of shared post-embryonic mutations between different cell-type lineages. We detected somatic mutations in different cell lineages covered in our dataset and identified a high degree of clonal expansion in myeloid cells. We used somatic mutations called from this method to construct neighbour-joining trees in macrophages and monocytes to infer that non-classical monocytes are the likely intermediary state between circulating monocytes and the majority of tissue resident macrophages in kidney cancer. We envisage that in the future, the use of spatial imaging techniques to visualise called mutations in expressed genes across a range of cell types will help to decipher the phylogenetic organisation of the multicellular TME.

An EMT meta-programme was defined through its expression by a subpopulation of tumour cells in each patient and shared by multiple ccRCC tumours in our study. The more abundant tumour cell populations and the use of methods to help circumvent challenging batch variations allowed us to uncover this previously unreported feature (*26–28*). Expression of the EMT program in ccRCC tumour cells was inversely correlated with that of the PT programme, an epithelial expression signature. Meanwhile, EMT^high^ tumour cells in ccRCC tended to localise to the tumour-normal interface, which is the leading and migration edge of a tumour. These findings, similar to those have been reported in the scRNA-seq study of head and neck cancer (*16*), reflect the defining feature of EMT: the loss of epithelial characteristic in cells in favour of promoting their migration and invasion abilities (*45*).

Analysing cell–cell interactions revealed the heterogeneity of predicted intercellular communications associated with different spatial localisations in the TME of ccRCC. Particularly, by linking ligands and target genes of interest using NicheNet, we found that IL1B, specifically expressed by a subset of tissue resident macrophage cells enriched at the tumour-normal interface (TR Mac.2), could potentially promote tumour cells undergoing EMT (Fig. 7D). Expression of IL1B has been reported to positively correlate with tumour stages of RCC (*46*) and is associated with worse prognosis of patients with RCC in patients recruited to The Cancer Genome Atlas. In addition, inhibition of IL1B in RCC has been shown to induce tumour regression in a syngeneic murine model of RCC (*47*). IL1B blockade was also shown to reduce incident lung cancer in patients with atherosclerosis (*48*), and its use is now being investigated in several clinical trials. In our data, we indicate that the underlying mechanism that results in the unfavourable role of IL1B in RCC acts through the promotion of EMT through macrophage derived IL1B signalling. Exploiting this pathway could be therapeutically useful.

## Supplementary materials

### Experimental model and subject details

Human kidney and tumour tissues were collected through studies approved by UK NHS research ethics committees. All adult kidneys samples, except PD44967 were collected from patients enrolled in the DIAMOND study; Evaluation of biomarkers in urological disease (NHS National Research Ethics Service reference 03/018). Tumour PD44967 was collected from a patient enrolled in Characterisation of the immunological and biological markers of Renal cancer progression (NHS National Research Ethics Service reference 16/WS/0039).

### Tissue sampling

Peripheral blood was sampled on the day of the surgery prior to removal of the kidney tumour and placed on ice. The surgical specimen was directly taken from the operating room to histopathology in order to minimise the warm ischaemia time. Biopsies were sampled by local pathologists to include (where available) multiregional tumour biopsies from 4 macroscopically disparate regions, the tumour-normal interface, normal kidney (distant to the tumour and close to cortico-medullary border), perinephric adipose tissue, and adrenal gland. The biopsy locations from the bivalved kidney were annotated. Tissue samples were divided and either placed on wet ice for immediate transfer for generation of single cell suspensions, or underwent formalin-free fixation for 24 hours in PAXgene Tissue FIX containers before being 20 transferred to PAXgene STABILIZER solution for storage at −20 °C.

### Generation of single cell suspensions

The fresh tissue samples were coarsely dissected using a single edged razor blade prior to digestion for 30 min at 37°C with agitation in a digestion solution containing 25μg/ml Liberase TM (Roche) and 50μg/ml DNase (Sigma) in RPMI (Gibco). Following incubation samples were transferred to a C tube (Miltenyi Biotec) and processed on a gentle MACS (Miltenyi Biotec) on programme spleen 4 and subsequently lung 2. The resulting suspension was passed through a 70μm cells strainer (Falcon), and washed with PBS. Percoll (Sigma-Aldrich) density separation was used both as a strategy to remove dead cells and cellular debris, and also to enrich stromal components of the TME, whilst still being permissive for a proportion of RCC cells themselves. We added the cell pellet to 44% Percoll in phosphate buffered saline (PBS) prior to centrifugation at 800G for 20 min. The supernatant was removed and the pellet suspended in PBS prior to centrifugation for 5 min at 800G. The concentration of enriched live cells was calculated after counting with a hemocytometer with trypan blue staining.

### Cell loading and 10x library preparation

Cells were loaded according to standard protocol of the Chromium single cell 5’mRNA kit with TCR library enrichment in order to capture approximately 14000 cells/chip position. All the following steps were performed according to the standard manufacturer protocol. Sequencing of libraries used either the Illumina HiSeq or NovaSeq systems.

### Initial processing of scRNA-seq data

After the conversion of CRAMs files into FASTQs using samtools (*49*), we used the 10X software package cellranger (version 2.1.1 and vdj) and the GRCh38 reference genome for processing the 5’ sequencing data. We used SoupX (*50*) to return an adjusted count matrix to account for ambient RNA contamination per channel using the adjustCounts() function. We then used DoubletFinder (*51*) to estimate the probability of a given droplet containing RNA from more than one cell. Given that our cell loading aimed to recover 14000 cells per lane, we assumed an 11% doublet formation rate.

### scRNA-seq merge and QC

Seurat V3’s (*52*) implementation of Reciprocal PCA (RPCA) was used to reduce the computational expense in merging the patient specific scRNA-seq data. Cells with greater than 30% mitochondrial content, or expression of fewer than 200 genes were excluded from further analysis. We used relatively permission thresholds to avoid removing renal epithelial cells that are known to have relatively high mitochondrial contents. We used standard clustering metrics and the expression of canonical marker genes to broadly classify cells into the principal cell subsets; T and NK cells, B and plasma cells, myeloid cells, endothelial cells, epithelial cells (non-cancerous), fibroblasts, and cancerous RCC cells. Cell clusters expressing implausible combinations of cell lineage specific marker genes were labelled as doublets and were excluded from further analysis.

### Cell type sub-clustering and annotation

We performed sub-clustering analysis of various cell compartments using the Seurat pipeline. Briefly, we first pulled out each cell compartment using the subset() function based on the broad classification of cells. We then used regularized negative binomial regression to normalize UMI counts using the SCTransform() function in Seurat, with the percentage of mitochondria genes being regressed out. Principal component analysis (PCA) was performed using the RunPCA() function based on highly variable features generated by using the VariableFeatures() function. For the PCA of T cell population, we excluded TCR encoding genes from the list of highly variable features so that to avoid clusters driven by the expression of different TCR genes. Batch correction was performed in each cell compartment using the RunHarmony() function implemented in the R package harmony, with the batch key (parameter ‘group.by.vars’) being set as patients and the assay (parameter ‘assay.use’) being set as ‘SCT’. Next, we performed nearest-neighbor graph construction, cluster determination and nonlinear dimensionality reduction using the FindNeighbors(), FindClusters() and RunUMAP() functions, respectively. The ‘reduction’ parameter in the FindNeighbors() and FindClusters() was set as ‘harmony’. DEGs of different clusters were extracted using the FindAllMarkers() function. Cell clusters expressing implausible combinations of cell lineage specific marker genes were labelled as doublets and were excluded from the analysis. Cell type annotation was based on the expression of canonical markers and DEGs in various clusters. The annotation of cell cycle phases in the T cell population was based on the previously reported phase specific genes (*17*).

### Pseudotime inference, TCR analysis

Single cell count data and associated metadata of CD8^+^ T cells was analysed using Monocle3 (https://github.com/cole-trapnell-lab/monocle3) after removal of cycling, gamma delta and MAIT cells. Pre-processing used the function preprocess_cds() with a dimensionality of 100, prior to alignment with ‘align_cds’ and batch correcting by individual sample. Dimension reduction used the function reduce_dimension(), prior to fitting the principal graph using ‘learn_graph’ and then ordering the cells using ‘order_cells’, all using the default parameters. To visualise the relationship of canonical marker genes of CD8^+^ T cell exhaustion we used the function plot_genes_in_pseudotime(). All such genes were found to be differentially expressed across the single cell trajectory using the function ‘graph_test’ at a *q* value of 0.

To demonstrate the differentiation properties of cells within clonotypes, we selected the most expanded clonotypes. For ease of interpretation we selected those clonotypes that contained at least 100 CD8^+^ T cells. The median, interquartile range, minimum, maximum values, and outlier values of psuedotime were plotted by clonotype, ordered by mean pseudotime values. The percentage maximum expansion was calculated from the region that contributed the maximum percentage of CD8^+^ T cells for each clonotype. The percentage of cells cycling in either the G1/S or G2/M phases were also calculated for each clonotype. We sought to quantify the degree of restriction of TCR clonotypes to a range of pseudotime values, by calculating the Wilcoxon test statistic for each clonally expanded CD8^+^ T cell clone (clones with more than one cell), compared to all of the other CD8+ T cells. To determine the likelihood of detecting expanded TCR clones in the blood as a function of pseudotime we computed the conditional density of detection of any cells with a given TCR in the blood, with psuedotime, for minimal clone sizes of 2, 4, 8, 16, 32, and 64 cells.

### Laser capture microdissection, library preparation, and low-input DNA sequencing

Laser capture microdissection and low-input DNA sequencing followed the protocol previously reported (*53*). Briefly, PAXgene fixed samples were subsequently embedded in paraffin using standard histological tissue processing. 16μm sections were cut, mounted onto PEN-membrane slides, and stained with Gill’s haematoxylin and eosin. Using the LCM (Leica LMD7), tumour regions were selected in order to perform focally exhaustive tumour sampling. The dissected cells were collected into separate wells in a 96-well plate. Tissue lysis was performed using Arcturus PicoPure Kit (Applied Biosystems).

Libraries were constructed using enzymatic fragmentation as described previously and subsequently submitted for whole-exome sequencing on the Illumina HiSeq X platform. Short insert (500bp) genomic libraries were constructed, flowcells prepared and 150 base pair paired-end sequencing clusters generated on the Illumina HiSeq X platform without PCR amplification. The average sequence coverage was 84X and 92X for tumour and normal dissection samples, respectively (table S3).

### Mutation calling from whole-exome sequencing

DNA sequencing reads were aligned to the GRCh 37d5 reference genome using the Burrows-Wheeler transform (BWA-MEM) (*54*). Single base somatic substitutions were called using an in-house version of CaVEMan v1.11.2 (Cancer Variants through Expectation Maximization, https://github.com/cancerit/CaVEMan). CaVEMan compares sequencing reads from tumour and matched normal samples and uses a naïve Bayesian model and expectation-maximization approach to calculate the probability of a somatic variant at each base. Small insertions and deletions (indels) were called using an in-house version of Pindel (github.com/cancerit/cgpPindel). Post-processing filters required that the following criteria were met to call a somatic substitution:

1. At least a third of the reads calling the variant had a base quality of 25 or higher.
2. If coverage of the mutant allele was less than 8, at least one mutant allele was detected in the first 2/3 of the read.
3. Less than 5% of the mutant alleles with base quality ≥ 15 were found in the matched normal.
4. Bidirectional reads reporting the mutant allele.
5. Not all mutant alleles reported in the second half of the read.
6. Mean mapping quality of the mutant allele reads was ≥ 21.
7. Mutation does not fall in a simple repeat or centromeric region.
8. Position does not fall within a germline insertion or deletion.
9. Variant is not reported by ≥ 3 reads in more than one percent of samples in a panel of approximately 400 unmatched normal samples.
10. A minimum 2 reads in each direction reporting the mutant allele.
11. At least 10-fold coverage at the mutant allele locus.
12. Minimum variant allele fraction 5%.
13. No insertion or deletion called within a read length (150bp) of the putative substitution.
14. No soft-clipped reads reporting the mutant allele.
15. Median BWA alignment score of the reads reporting the mutant allele ≥ 140. The following variants were flagged for additional inspection for potential artefacts, germline contamination or index-jumping event:
16. Any mutant allele reported within 150bp of another variant.
17. Mutant allele reported in >1% of the matched normal reads.
18. The median alignment score of reads that support a mutation should be greater than or equal to 140 (ASMD ≥ 140)
19. Fewer than half of the reads should be clipped (CLPM = 0).

We then tested for true presence or absence of the somatic variants that passed the above flags using an approach previously described (*55*). Briefly, counts were re-calculated using AlleleCounter (https://github.com/cancerit/alleleCount) across all the samples in this study. For each patient, the non-tumour samples in this study not belonging to that patient were used as a reference to obtain the locus-specific error rate. To minimize the false positive rate, the presence of the variant in the sample was accepted if the multiple-testing corrected p-value was less than 0.001. The ascatNGS (*56*) algorithm was used to estimate tumour purity and ploidy and to construct copy number profiles. A penalty of 200 was used with the prior knowledge that copy number events in RCC tended to be either arm or chromosome level.

### DNA mutational clustering

Mutations were clustered using a Bayesian Dirichlet based algorithm as described previously (*57*). Briefly, the expected number of reads for a given mutation if present in one allelic copy of 100% of tumour cells may be estimated based upon the ASCAT derived tumour cell fraction, the copy number at that locus and the total read-depth. The fraction of cells carrying a given mutation is modelled by a Dirichlet process with an adjustment for the decreased sensitivity in identifying mutations in lower tumour fractions. Mutations were thus assigned to clusters according to the calculated fraction of clonality. The hierarchical ordering of these clusters was determined by applying the pigeonhole principle.

### *De Novo* Mutation Calling from scRNA-seq Data

The code for this method is available at https://github.com/ThomasJamesMitchell/deSCeRNAMut. The steps are described below:

1. Initial variant calling In order to call cell specific mutations, indexed BAM files from the cellranger pipeline were first split into cell specific BAM files and were indexed using samtools (*49*). Mutations were initially called using bcftools mpileup. The choice of mutation caller was primarily influenced by the need for high sensitivity calls of variants with few supporting reads (*58*). Unsurprisingly, a huge number of mutations were called - with between 800,000 and 4,000,000 mutations called per patient. To facilitate more efficient downstream filtering of putative mutations, we perform the first filter step at this point:

- Removal of singlet variants only called in a single cell as it will be challenging to accurately determine whether these mutations are real or artefact.
- Removal of variants that are shared between the main cell lineages of T and NK cells, B and plasma cells, myeloid cells, endothelial cells, epithelial cells (non-cancerous), fibroblasts, and cancerous RCC cells. The vast majority of somatic mutations are acquired post embryonic differentiation, and therefore any true degree of sharing is implausible. After these steps, we are left with between 40,000 and 300,000 mutations per patient. We have generated a list of putative variant sites, but we are unaware how many variants may have been missed at each loci, and we have no information regarding reference calls at those loci. We therefore run alleleCount (https://github.com/cancerit/alleleCount) to generate count tables of each base for all cells at every putative patient-specific loci.
2. Collation and annotation of counts Reference and variant counts were collated for all of the loci called above to create a sparse matrix of counts for all cells. In the absence of copy number variants, if an autosomal chromosome harbours a true mutation, one expects an approximately equal number of reference and variant calls. The exception is for genes that exhibit a high degree of allelic specific expression, or that typically transcribe a particular allele in concentrated bursts. Alternatively, a high ratio of reference to variant counts in a cell base may imply artefact associated with high depth sequencing/ poorly mapped regions. A binomial filter (*p* < 0.05) was therefore applied in each cell, with calls ignored in future analyses if there are significantly higher reference than variant counts. Each genomic loci was annotated using ANNOVAR (*59*) and the trinucleotide context of the variant. The number of cells containing either the reference or variant base were collated for:

- The cell lineage with the greatest number of mutations.
- All of the other cell lineages.
- The TCR clonotype with the greatest number of mutations.
- All other TCR clonotypes Fisher’s exact test was used to compute whether there are proportionally greater numbers of mutations in the cell lineage/ clonotype with the greatest number of mutations. An enrichment factor was also calculated for each mutation that represents the multiple of the increased prevalence in the predominant cell type compared to all others.
3. Final filter We applied the following thresholds to filter all possible mutations

- Fisher’s exact significance of enrichment by cell lineage, *p* < 0.0001 with proportionally at least 5 times greater mutations in the most enriched lineage.
- Absence of any known single nucleotide polymorphisms from either ExAC or dbSNP.
- No shared mutations between patients
- Adequate coverage with at least 5 cells with variant base from the mutated cell lineage and at least 20 cells with reference base from the reference population We then examined the trinucleotide context of called mutations after this filtering step. Note is made of high levels of mutations that are otherwise unexplained from published catalogues of mutational signatures (particularly in a GCN>GGN and GTN>GGN context). By separating the trinucleotide context into the positive versus negative transcribed strands, we see differences that are otherwise unexplained by DNA derived mutational signatures, implying artefact either through library prep, sequencing, or RNA editing. The striking strand bias cannot be accounted for by known mutational processes. Given the disparity between transcribed strands, mutations that have arisen with a highly biased context are removed (binomial filter, *p* < 0.005). We finally removed all mutations that are clustered within 4 bases in a given patient, to yield the final mutation calls.
4. Benchmarking Data by whole-exome sequencing Multi-regional whole exome sequencing data has been processed for tumour tissue adjacent to the regions that have undergone single cell RNA sequencing. The exonic mutations may therefore be used as a benchmark to determine the precision and sensitivity of the single cell mutation calling method above. To provide a fair comparison between single cell RNA and bulk exonic DNA mutation calls, and to account for differences in coverage between the methods, we also examine whether there is evidence of a given mutation using the reciprocal technology by performing a pileup at that mutation locus. We can therefore classify mutations called using the above pipeline as:

- True positive - The mutation has been called in both in the scRNA-seq pipeline and CaVEMan.
- True positive, pileup only - The mutation has been called in the scRNA-seq pipeline, and there is evidence of the mutation in exome sequencing from tumour regions, with no mutations in the normal sample BAM files. The most common reasons for these mutations not being called by CaVEMan is low coverage or the mutation being called in mitochondrial DNA.
- False positive - The mutation has been called in the scRNA-seq pipeline, but there are fewer than 5 supporting reads for the variant base, and more than 20 reads for the reference base in the exome data.
- False negative - The mutation has been called by CaVEMan from the exome data, and has not been called from the scRNA-seq data, despite there being adequate coverage of at least 5 cells with the variant and at least 20 cells with the reference base.
- Indeterminate - The mutation has been called by the scRNA-seq pipeline, but there is not sufficient depth in the exome data to corroborate the call. Note that it is possible that some of the false positive results may be real mutations that simply have not been captured spatially as adjacent tissue was sequenced. Overall, this scenario is unlikely as the majority of mutations are clonal and present throughout the tumour.
5. Benchmarking Data by clonotype

In adult tumours, one expects a high proportion of somatic mutations in expanded CD8^+^ T cells to have been acquired post thymic selection. Most called mutations should therefore be restricted to a single T cell receptor clonotype. By using identical metrics to those used to select mutations across all cell types, we examined the proportion of CD8^+^ T cell mutations that are restricted to a single clonotype. Again, in order to call a mutation, we use thresholds requiring at least 5 cells with the variant in the most prevalent clonotype, with a least 20 cells covering the reference allele in the other clonotypes

### Inferring copy number variations based on scRNA-seq data

To effectively distinguish malignant and non-malignant cells, we inferred the large-scale chromosomal CNVs of single cells based on scRNA-seq data using the tool InferCNV (https://github.com/broadinstitute/inferCNV) with default parameters. Briefly, InferCNV first orders genes according to their genomic positions (first from chromosome 1 to X and then by gene start position) and then uses a previously described sliding-average strategy to normalize gene expression levels in genomic windows with a fixed length. Multiple putative non-malignant cells are chosen as the reference to further denoise the CNV result. In our analysis, we chose epithelial cells (including both PT and non-PT cells), endothelial cells and fibroblasts as the reference cell types to define a baseline in inferring CNVs.

### Cell subtype abundance in different tissues

To explore the potential enrichments of cell subtypes in different tissues, we compared the observed and expected number of cells in each subtype across different tissues. The ratio of observation to expectation (R_O/E_) was calculated as follows:

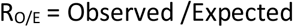

where the expected number of cells were calculated based on the Chi-square test. In this analysis, we excluded cells from the adrenal metastasis and tumour thrombus because we only captured cells from these two tissues from single patients. A specific cluster was considered as being enriched in a specific tissue if Ro/e > 1.

### Correlation between spatial, somatic RCC evolution and TCR clonotype evolution

Tree structures relating to somatic ITH, spatial localisation of the tissue samples, and CD8^+^ clonotype enrichment for each region sampled were generated. The distance matrix relating regions to their somatic ITH was generated using pairwise distances from the mutational cluster output from the Bayesian Dirichlet based algorithm from the WES data for each of the (clonally) derived LCM samples. The spatial localisation distance matrix was calculated from the pairwise distances from tree structures determined either by:

1. The approximate absolute distance between LCM biopsies: This metric is not meaningful for normal tissue samples, particularly for peripheral blood and therefore the normal samples were excluded using this absolute distance metric.
2. A categorical distance: The first level equates to adjacent LCM biopsies, whose centres lie approximately 0.2mm apart. The second level for LCM biopsies taken from the same histologically mounted section, approximately 2mm distant. The third level relates to biopsies from small macroscopically separate biopsies, separated by approximately 6mm. The fourth level relates to macroscopic tumour biopsies taken approximately 30mm apart. The fifth level encompasses all of the adjacent normal tissue samples.

The Euclidian CD8^+^ T cell clonotype distance matrix was calculated using the relative expansions of the CD8^+^ clonotypes for each region sampled. Regions were removed where there was incomplete data – for instance if there were no viable cells in the single cell sequencing data. However, any regions where there was overlapping data, for instance multiple WES data from adjacent LCM cuts relating to a single region for single cell RNA sequencing were all included.

The pairwise correlation between the above distance matrices was computed using the Mantel test. A paired Wilcoxon test was used to determine whether somatic ITH or spatial localisation correlated with CD8^+^ clonotypic heterogeneity.

### Gene set enrichment analysis and gene signature scoring in macrophage population

We performed gene set variation analysis among macrophage subsets using the GSVA R package. The gene sets we used were the C2 collection (curated gene sets) downloaded from the MsigDB database (https://www.gsea-msigdb.org/gsea/msigdb). The differences in activities pathways between clusters were calculated using the Limma R package. Significantly disturbed pathways were identified with Benjamini-Hochberg–corrected P value of <0.01. Some representative pathways that related to tumour progression, immune response and regulation were selected to make a heatmap. We investigated the phenotypes of different macrophage subsets by scoring them based on four previously reported gene signatures, including M1 and M2 polarization (*18*), signature of myeloid-derived suppressor cells (MDSC) (*60*), and signatures of angiogenesis and phagocytosis (*33*)

### RNA velocity analysis

We conducted RNA velocity analysis using velocyto (*61*). We first ran the command line ‘velocyto run10x’ to annotate spliced and unspliced reads using the cellranger output (the BAM file) as the input, generating loom files for each cellranger output. We then merged these loom files and pre-processed the velocity data using the scVelo python package (*62*). We projected the velocity information onto pre-generated UMAP and visualized the results using the function scvelo.pl.velocity_embedding_grid().

### Similarity analysis of myeloid clusters

To compare the similarities of myeloid clusters to the previously published data (*33*), we trained a logistic regression model using elastic net regularization as previously described (*25*). The previous kidney cancer data were obtained from Gene Expression Omnibus (GEO: GSE154763) and were used as training data.

### Lineage tracing using scRNA-seq called somatic mutations

Based on the somatic mutations called from scRNA-seq data, we constructed a neighbor-joining tree to elucidate the relationship of different monocyte and macrophage subtypes (the low-quality cluster was excluded). Since our somatic mutations were called from gene expression data, we realized that the expression levels of genes may impact on the detection of mutations in different clusters, thus potentially making cell subtypes with more similar expression profiles cluster closer while those with less similar expression profiles segregate farther in the tree structure. To mitigate this, we excluded mutations that were detected in the top 100 DEGs of every cluster from the tree construction process. Based on the remaining mutations, we created a mutation matrix (mutation × subtype) considering whether a specific mutation appears in specific subtypes or not. Next, we calculated the binary distance between any two cell subtypes based on the mutation matrix and constructed the neighbor-joining tree using the ‘NJ’ function in the R package ‘phangorn’. A bootstrapping analysis was performed using the ‘boot.phylo’ function implemented in the R package ‘ape’, with the number of bootstrap replicates being set as 100. The final tree structure was displayed using the ‘plotBS’ function in the R package ‘phangorn’.

### Deciphering intra-tumour expression programmes and meta-programmes

To explore underlying intra-tumour expression signatures of tumour cell population in RCC, we applied non-negative factorization (implemented in the R NMF package) to the tumour cells in ten patients (PD44714 and PD47172 were excluded from this analysis because they were histologically evaluated as benign and oncocytoma). Briefly, for each tumour, we first normalized the expression counts using Seurat NormalizeData() function with default parameter settings. We selected highly variable genes (HVGs) using Seurat FindVariableFeatures() function and only focused on the 2000 HVGs in downstream analysis Then, we performed center-scale for HVSs using Seurat ScaleData() function with the percentage of mitochondria genes being regressed out, and replaced all negative values in the expression matrix by zero. The top 10 ranked co-expressed gene modules in each tumour sample were dissected by using the nmf() function in the NMF package. For each gene module, we extracted the top 50 genes with the highest weight and used them to define a specific intra-tumour expression programme. Finally, we only included those expression programmes with standard deviations larger than 0.2 among tumours cells, thus generating 3 to 6 intra-tumour expression programmes in the 10 tumours.

To investigate if some intra-tumour expression programmes were actually shared by multiple tumours, we applied a clustering analysis to all programmes based on the pair-wised Jaccard index calculated as follows, where A and B represent two intra-tumour programmes.

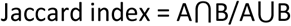

We defined those intra-tumour programmes shared by multiple tumours as meta-programmes. Genes that are shared by at least 50% tumours with a specific meta-programme were used to define the meta-programme except for the cell cycle programme, which is only shared by two tumours and thus we used genes shared by the two tumours to define the cell cycle programme.

### Integrating and analyzing tumour cells from different patients

To mitigate the effect brought by the strong inter-tumour heterogeneity in integration, we used the Seurat scRNA-seq integration pipeline to integrate tumour cells from 10 patients (PD44714 and PD47172 were excluded from this analysis because they were histologically evaluated as benign and oncocytoma). Briefly, for each tumour, we first used regularized negative binomial regression to normalize UMI counts based on the SCTransform() function in Seurat with the percentage of mitochondria genes being regressed out. The pre-processed individual objects were then added to a list, based on which we further performed selection of integration features using the SelectIntegrationFeatures() function with the number of features being set as 3000. We next performed integration preparation using the PrepSCTIntegration() function and found the integration anchors using the FindIntegrationAnchors() function with the normalization method being set as ‘SCT’ and the ‘k.filter’ parameter being set as 50. Finally, these objects were integrated by using the IntegrateData() function. Based on this integrated object of tumour cells, we further performed downstream analyses including clustering and differentially expressed gene analysis. Gene signature scores of the six identified meta-programmes were calculated with the AddModuleScore() function using featured genes in these programmes.

### TCGA data and prognosis analysis

We used TCGA expression and prognostic data to calculate meta-programme scores and investigate how the meta-programmes correlate with survival of patients with ccRCC. We processed the gene expression matrix by log-transforming and centralizing. Gene scores of each meta-programme were calculated as the average expression of genes in the specific programme. TCGA samples with records of age, gender, stage, survival data and tumour purity information were further used for survival analysis. For the expression of each meta-programme, patient cohorts were grouped into high and low groups by the optimal cut point determined using the cutp() function documented in the survMisc R package. We performed multivariate analyses using the Cox proportional hazards model (coxph() function in the survival R package) to correct clinical covariates including age, gender, tumour stage and tumour purity for all survival analyses in our study. Kaplan-Meier survival curves were plotted to show differences in survival time using the ggsurvplot() function in the survminer R package.

### Cell-cell interaction analysis

We inferred putative cell-cell interactions based on the expression of ligand-receptor pairs on different types of cells using CellPhonDB (*39, 40*). We investigated cell-cell interactions occurring between any two of the 12 major cell types in our study, and compared the numbers and types of interactions among the normal kidney, tumour-normal interface and tumour core. Next, we conducted a deeper analysis of cell-cell interaction by linking ligands expression on one cell type to some target genes of interest expressing another cell type using NicheNet (*41*). This analysis uses public databases (KEGG, ENCODE, PhoshoSite) to track downstream effectors such as transcription factors and receptor’s target in the provided dataset. Specifically, we were interested in what ligands from non-malignant cells in the TME can potentially trigger EMT programme in tumour cells, thus considering the gene list of deciphered EMT meta-programme as the target genes. Genes were considered as expressed when they have non-zero values in at least 5% of the cells in a specific cell type.

### Data and software availability

The accession number for the genome sequence data reported in this paper is European Genome-Phenome Archive: EGAD00001008029 for whole-exome sequencing data, and EGAD00001008030 for the single cell RNA sequencing data. Code and pipeline for deSCeRNAmut is available at https://github.com/ThomasJamesMitchell/deSCeRNAMut.

---

**Fig. S1.**
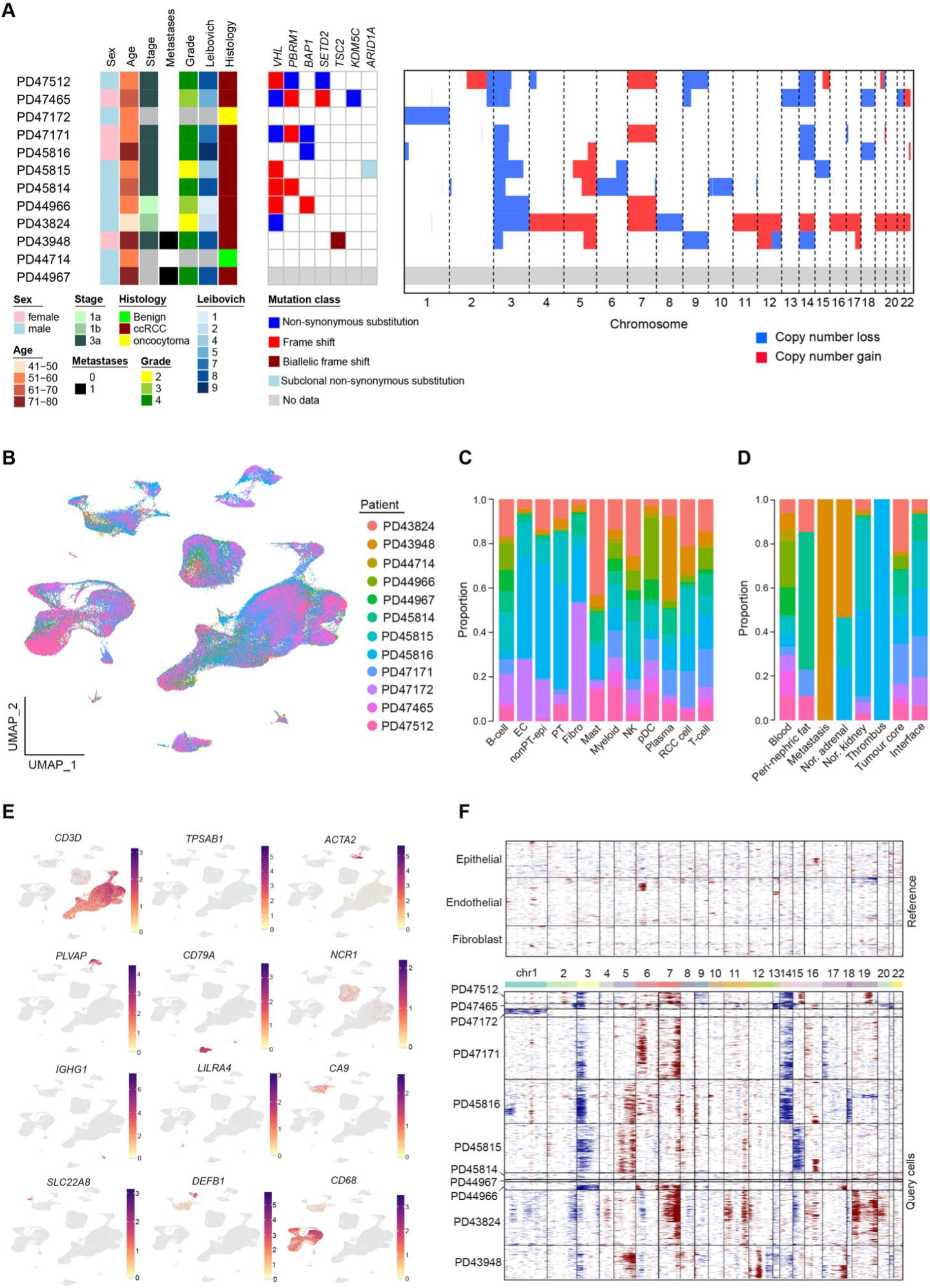
Basic information of the study cohort and the data. (A) Heatmap illustrating the clinical features (left panel) and the genomic landscape (middle panel) and copy number profiles (right panel) of the tumours sequenced. (B) UMAP, (C) cell type, and (D) sampled region depicting the interpatient variability of scRNA-seq results. (E) UMAP showing marker gene expression for all cells. (F) Copy number inference based on scRNA-seq data.

**Fig. S2.**
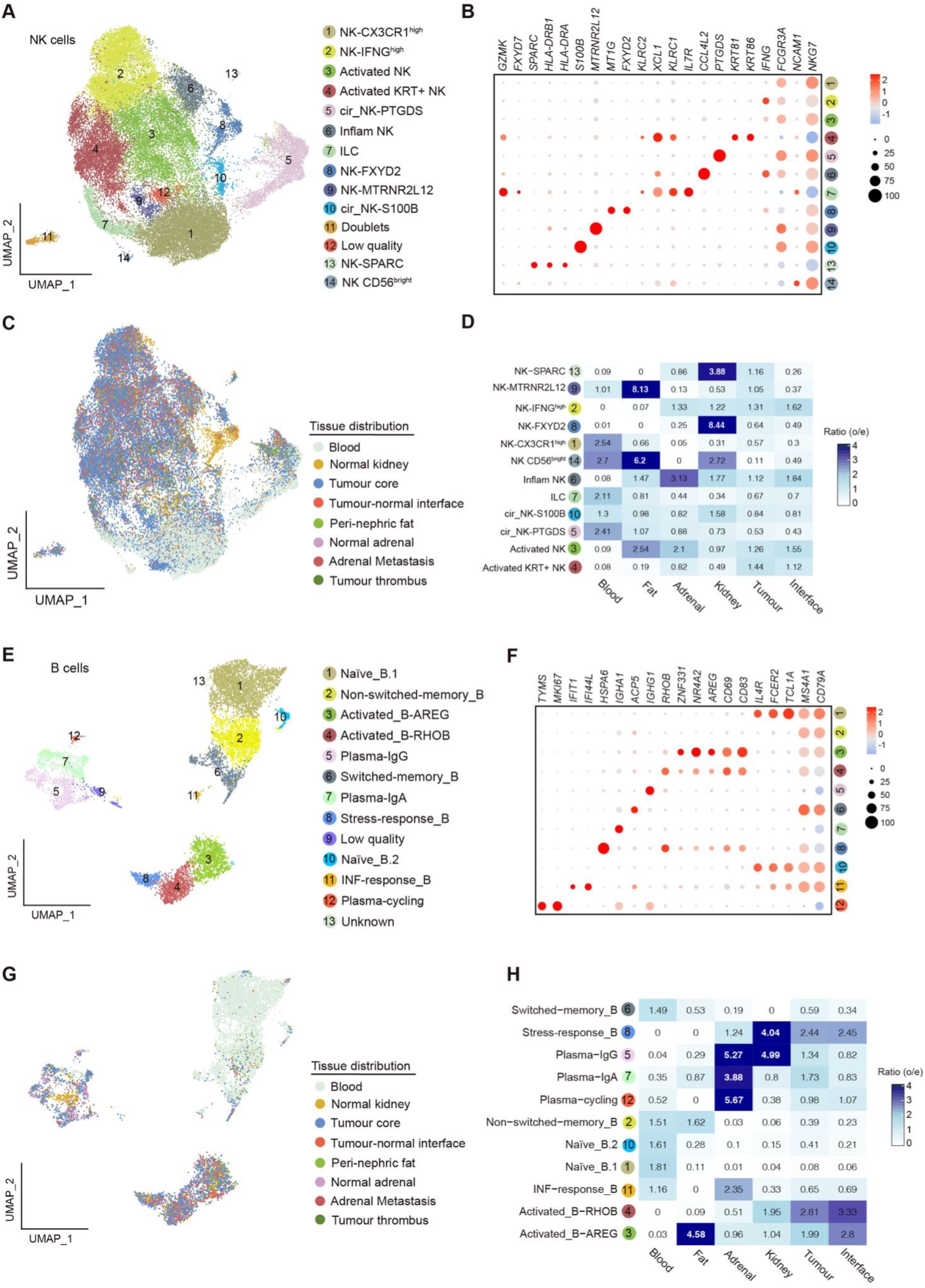
Spatial and transcriptomic heterogeneity of NK and B cell compartments. (A&E) UMAPs, (B&F) dotplots of marker genes, (C&G) tissue distribution, and (D&H) regional enrichment of NK and B cells.

**Fig. S3.**
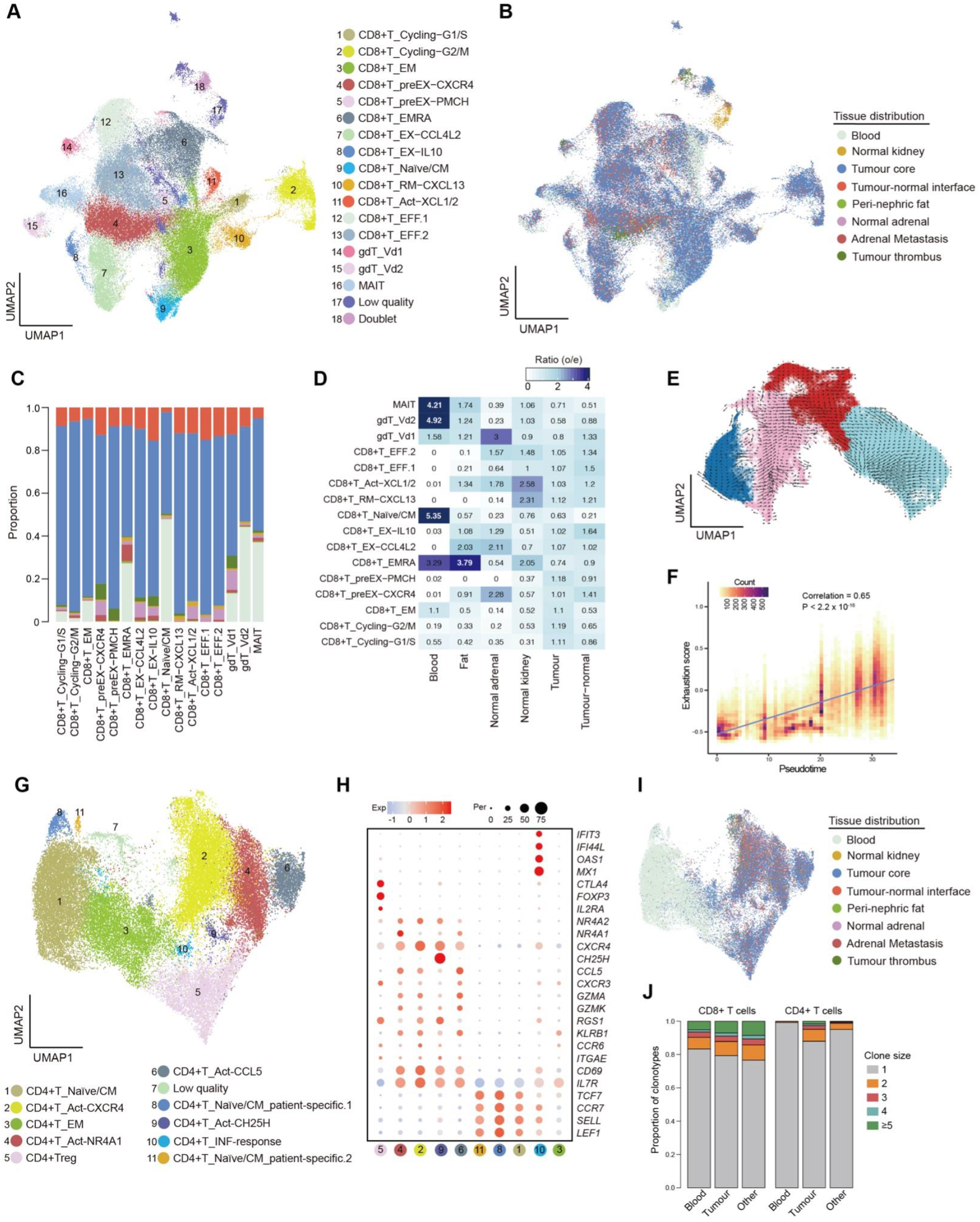
Spatial and transcriptomic heterogeneity of T cell compartment. (A) UMAP of the sub-clustering result, (B) UMAP and (C) barplot showing tissue distribution, and (D) regional enrichment of CD8+ T cell population. (E) UMAP showing the RNA velocity result and (F) exhaustion score across the pseudotime trajectory of CD8+ T cells. (G) UMAP of the sub-clustering result, (H) dotplot showing marker gene expression, and (I) UMAP of tissue distribution of CD4+ T cells. (J) Barplot showing the comparison of TCR clonal expansion between CD8+ and CD4+ T cells, breaking down into three different locations: blood, tumour and other regions.

**Fig. S4.**
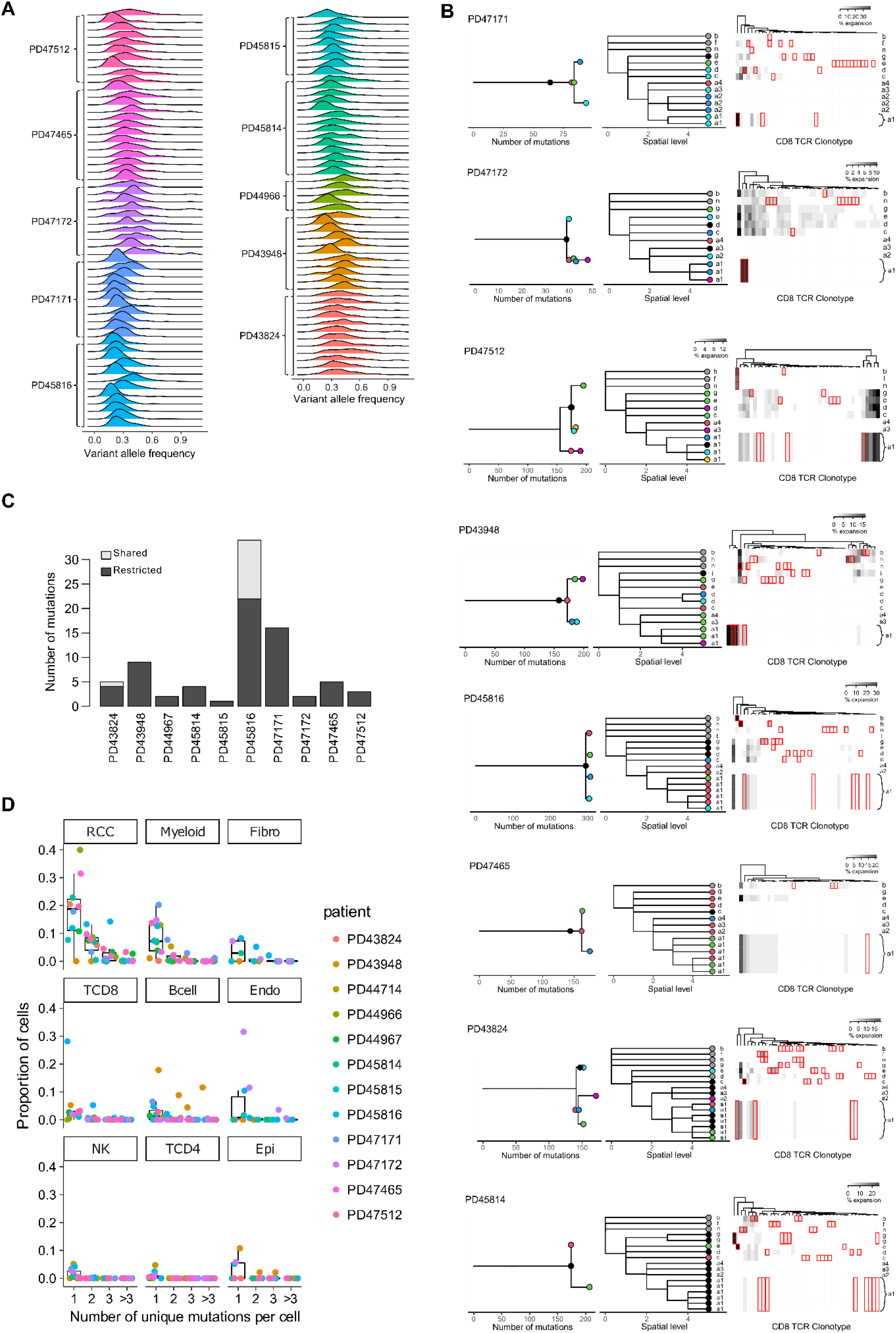
Somatic mutation analysis and the relationship with TCR heterogeneity. (A) Density plots showing the VAF distribution of mutations called from WES data from LCM biopsies of tumour samples. (B) Comparison of WES derived phylogenies (left) with geographic location (centre) and CD8+ TCR clonotype expansion (right) for each patient with matching data. Colours reference somatic clones to spatial localisation. Each column in the right panel represents a TCR clonotype, those with significant regional enrichment are highlighted in red. a, c, d, and e represents four different regions of the tumour core; g, tumour-normal interface; f, perinephric fat; n, normal kidney; b, peripheral blood; h, normal adrenal gland; i, adrenal metastasis; t, thrombus. (C) Bar chart showing benchmarking results of the number of called mutations in CD8+ T cells derived from scRNA-seq data that are restricted to individual TCR clonotypes. (D) Comparison of the proportion of cells with one, two, three, and more than three mutations across the major cell types.

**Fig. S5.**
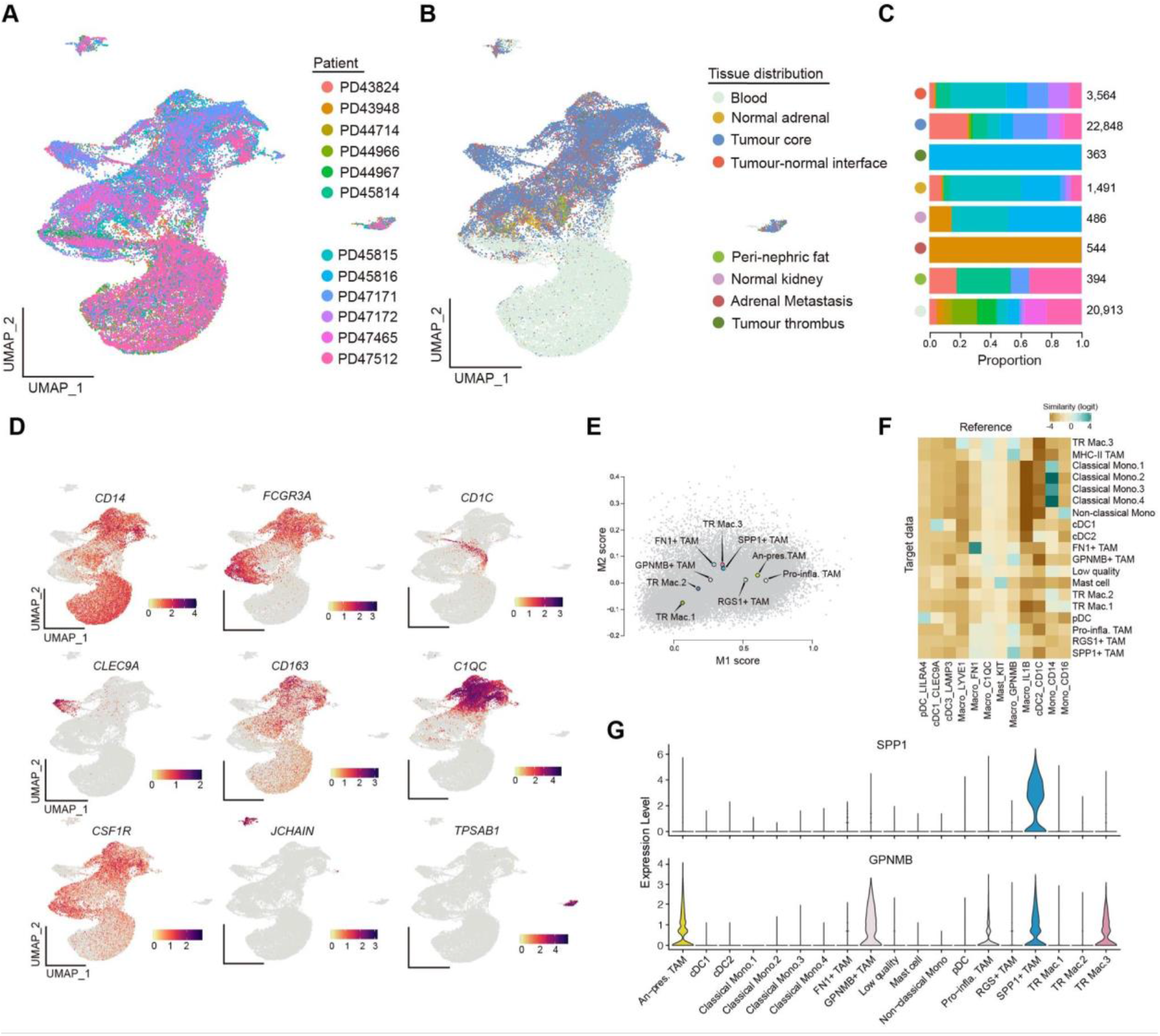
Sub-clustering analysis of myeloid cell compartment. (A) UMAP depicting interpatient variation, (B) tissue distribution, and (C) patient contribution to the different regions sampled. (D) UMAPs showing marker gene expression in myeloid cells. (E) Scatter plot of M1 versus M2 polarisation for all of the macrophages showing individual cells (grey), and mean values for each subtype (coloured). (F) Heatmap plotting the similarity of myeloid clusters derived in our study with those published in ref. 33. (G) Violin plots showing expression levels of SPP1 and GPNMB across all myeloid cell subtypes.

**Fig. S6.**
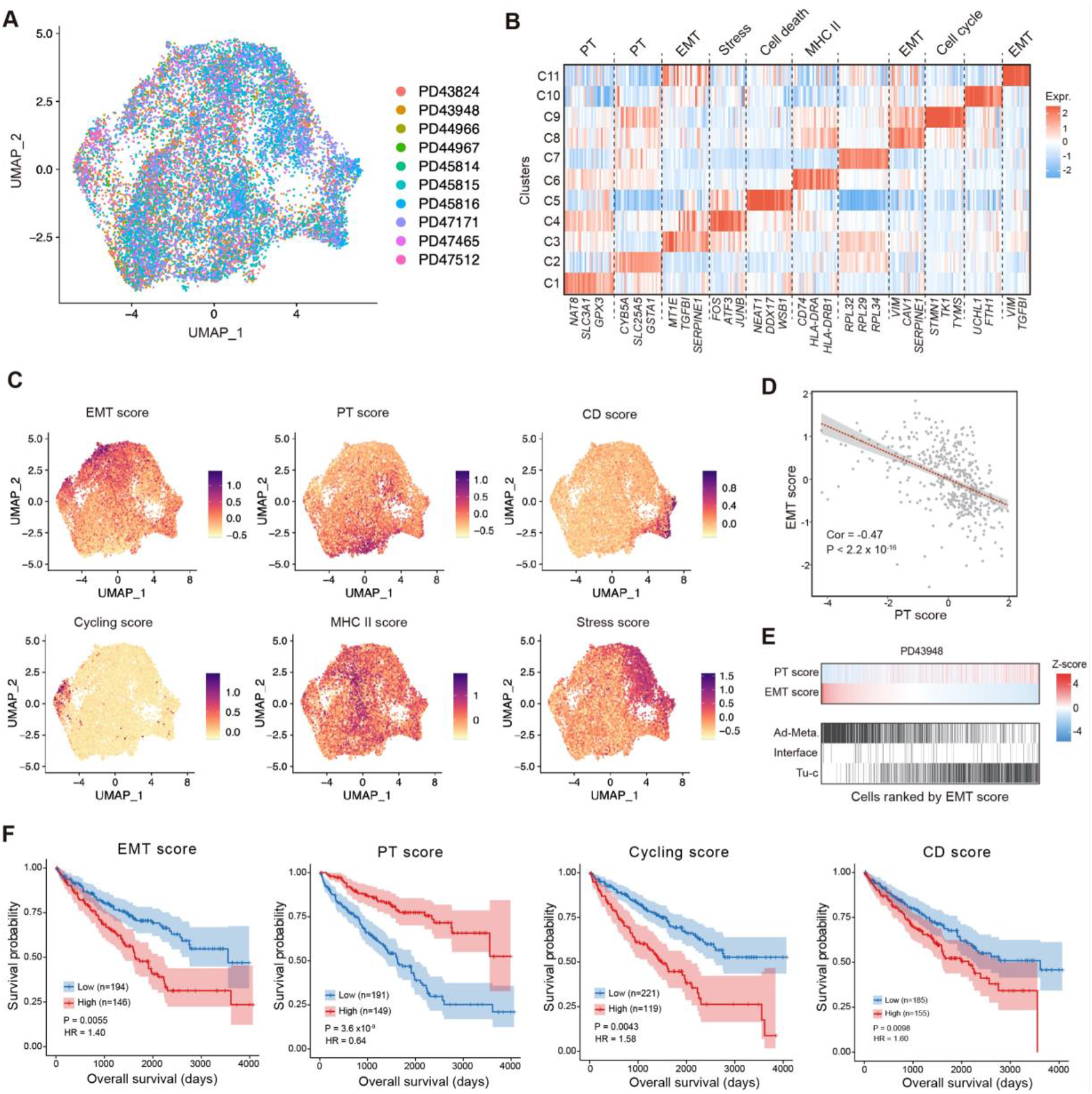
Tumour cell expression programmes. (A) UMAP showing the patients that RCC cells derived from. (B) Heatmap depicting the top DEG expression and assignment to meta-programmes for RCC cell clusters. (C) UMAPs showing the relative expression of each meta-programme for all RCC cells. (D) Correlation of EMT versus PT scores from bulk RNA sequencing data of the TCGA. (E) Cells from patient donors PD43948, ranked by decreasing EMT score with corresponding PT score and cell location. Tu-c, tumour core. (F) Survival probability of patients according to stratification of bulk RNA sequencing from TCGA with single-cell derived meta-programmes. HR, hazard ratio.

## Supplementary Tables

**Table S1. Clinicopathological information**

Table_S1.xlsx

**Table S2. Information of scRNA-seq**

Table_S2.xls

**Table S3. Information of LCM samples and WES**

Table_S3.xlsx

**Table S4. Top differentially expressed genes in the sub-clustering analysis of different cell lineages**

Table_S4.xlsx

**Table S5. Information of expression programmes**

Table_S5.xlsx

## AUTHOR CONTRIBUTIONS

Conceptualization, T.J.M., R.L.; Methodology, T.J.M., R.L.; Software, R.L., T.J.M.; Formal Analysis, R.L., T.J.M.; Investigation, R.L., J.R.F, K.W.L., G.S.B., L.M., J.B.N., L.B., E.S.F., Y.H., A.Y.W., M.G.B.T., S.B., M.D.Y; Resources, J.B.N., A.R.J.L., M.D.Y., T.R.W.O., T.M.B., J.N.A., T.A., A.C.P.R., V.G., M.G.B.T., G.D.S., M.R.C.; Project administration, T.J.M., S.J.W., K.B.M., M.R.C., P.J.C., S.A.T.; Supervision; P.J.C., S.A.T.; Funding acquisition, T.J.M., S.A.T., P.J.C.; Writing – original draft, R.L, T.J.M.; Writing – review and editing, R.L., T.J.M, S.T., M.R.C., S.B., M.Y., S.W., G.D.S., J.B.N., M.G.B.T.

## ACKNOWLEDGEMENTS

This work was supported by Cancer Research UK/Royal College of Surgeons Clinician Scientist Fellowship (T.J.M: C63474/A27176), British Heart Foundation (R.L), the National Institute of Health Research (NIHR) Cambridge Biomedical Research Centre and the NIHR Blood and Transplant Research Unit (J.R.F. and M.R.C), Kidney Research UK Clinical PhD Fellowship (K.W.L.: TF_013_20171124), Wellcome Science Strategic Award for the Human Cell Atlas (G.S.B), Medical Research Council Human Cell Atlas Research Grant (M.R.C.: MR/S035842/1), Cancer Research UK Cambridge Centre (A.W.: C9685/A25177), Kidney Cancer UK and Facingup2Kidney (M.G.B.T). The Wellcome Sanger Institute is supported by core funding from the Wellcome Trust (206194).

Tissue samples were acquired as part of the DIAMOND study “Evaluation of biomarkers in urological disease” - NHS National Research Ethics Service reference 03/018, whose infrastructure is part-funded by the NIHR Cambridge Biomedical Research Centre (BRC-1215-20014) and CRUK Cambridge Centre Urological Malignancies programme (Cancer Research UK Major Centre Award C9685/A25117). The views expressed are those of the authors and not necessarily those of the NIHR or the Department of Health and Social Care. Tissue and blood processing was carried out in the Clatworthy Lab, based in the University of Cambridge Molecular Immunity Unit in the MRC Laboratory of Molecular Biology and we are grateful for the use of their core facilities. We acknowledge the assistance from the CASM core team at the Wellcome Sanger Institute, in particular Laura O’Neil and Kirsty Roberts for assistance with tissue handling and library preparation.

